# Force propagation between epithelial cells depends on active coupling and mechano-structural polarization

**DOI:** 10.1101/2022.06.01.494332

**Authors:** Artur Ruppel, Dennis Wörthmüller, Vladimir Misiak, Manasi Kelkar, Irène Wang, Philippe Moreau, Adrien Méry, Jean Révilloud, Guillaume Charras, Giovanni Cappello, Thomas Boudou, Ulrich S. Schwarz, Martial Balland

**Affiliations:** Laboratoire Interdisciplinaire de Physique, Grenoble Alpes University, Saint Martin d’Heres, France; Institute for Theoretical Physics, Heidelberg University, Heidelberg, Germany; BioQuant–Center for Quantitative Biology, Heidelberg University, Heidelberg, Germany; London Centre for Nanotechnology, University College London, London, United Kingdom; Department of Cell and Developmental Biology, University College London, London, United Kingdom; Institute for the Physics of Living Systems, University College London, London, United Kingdom

## Abstract

Cell-generated forces play a major role in coordinating the large-scale behavior of cell assemblies, in particular during development, wound healing and cancer. Mechanical signals propagate faster than biochemical signals, but can have similar effects, especially in epithelial tissues with strong cell-cell adhesion. However, a quantitative description of the transmission chain from force generation in a sender cell, force propagation across cell-cell boundaries, and the concomitant response of receiver cells is missing. For a quantitative analysis of this important situation, here we propose a minimal model system of two epithelial cells on an H-pattern (“cell doublet”). After optogenetically activating RhoA, a major regulator of cell contractility, in the sender cell, we measure the mechanical response of the receiver cell by traction force and monolayer stress microscopies. In general, we find that the receiver cells shows an active response so that the cell doublet forms a coherent unit. However, force propagation and response of the receiver cell also strongly depends on the mechano-structural polarization in the cell assembly, which is controlled by cell-matrix adhesion to the adhesive micropattern. We find that the response of the receiver cell is stronger when the mechano-structural polarization axis is oriented perpendicular to the direction of force propagation, reminiscent of the Poisson effect in passive materials. We finally show that the same effects are at work in small tissues. Our work demonstrates that cellular organization and active mechanical response of a tissue is key to maintain signal strength and leads to the emergence of elasticity, which means that signals are not dissipated like in a viscous system, but can propagate over large distances.

## Introduction

Cell-generated forces are essential for tissue morphodynamics, when eukaryotic cells change number, shape and positions to build a multi-cellular tissue. Tissue morphogenesis is a dominant process during development, but also occurs in adult physiology and disease, in particular during wound healing and cancer, respectively. In addition to driving cell shape change and movement, force-producing processes allow cells to probe the mechanical and geometrical properties of their environment [1, 2], feeding back on to major cellular processes, such as differentiation [3, 4, 5, 6], fate [7, 8, 9] or migration [10, 11, 12]. Generation of contractile force is an universal property of mammalian cells due to the ubiquitous expression of non-muscle myosin II [13]. It is less clear, however, how this force is propagated through tissue and how long-ranged its effects are. Fast and long-ranged propagation of mechanical force seems to be essential during development, when morphogenesis has to be coordinated across the embryo [14, 15]. For example, the onset of migration of neural crest cells in Xenopus appears controlled by the stiffening of the underlying mesoderm resulting from axis elongation [16]. An example of a more mature tissue is the epithelium of the ju-venile oesophagus in mice, whose transition from growth to homeostasis is mediated by the mechanotransduction of progressively increasing mechanical strain at the organ level [17].

Despite these interesting observations for development, it is not clear how force is propagated across tissues in general and whether propagation is passive or sustained by mechanochemical feedback loops. Force propagation across tissues suffers from the same challenge as any other information propagation through a passive medium. Whether it be an electrical signals transmitted through a telegraph line or an action potential originating in the soma of a neuron, the signal typically attenuates with distance until it becomes indistinguishable from noise [18]. The main measure to counteract such attenuation are active processes that restore signal strength, like the opening of voltage-gated ion channels along the axon for action potentials. In addition to electrical currents, mechanical waves have also been observed to propagate along lengths several orders of magnitude larger than the cell-size in confined epithelial tissues [19]. These waves require active cellular behaviours such as contractility and F-actin polymerization to propagate, suggesting that cells actively respond to external forces to maintain the strength of the signal as it propagates through the tissue [20, 21, 22]. Furthermore, it has been shown that passive cells in an epithelial tissue act as obstacle for mechanical wave propagation [23]. Despite these studies, our knowledge of force propagation remains largely qualitative because of the lack of a model system that allows for precise spatiotemporal control of force generation and quantitative characterization of the propagation of the mechanical signal across intercellular junctions. As a result, we know little of how far force signals can propagate from their origin or whether signal propagation efficiency depends on tissue organization. Indeed, in some tissues, such as the hydra ectoderm, stress fibers within the cells of the ectoderm form a nematic system [24]. This high degree of alignment of force-generating subcellular structures suggests that tissues may display anisotropic propagation of stresses.

Here, we introduce such a sought-after minimal biophysical system for force propagation in epithelia, consisting of two interacting cells in which force generation is controlled by an optogenetic actuator of contractility and force propagation is quantitatively monitored using traction and monolayer force microscopies. To place the two cells next to each other with a stable cell-cell boundary, we make use of adhesive micropatterning [25, 26]. Moreover, the adhesive micropatterning allows us to control the aspect ratio of the cells and the structural organization of their cytoskeleton. Using this system, we show that intercellular force propagation is an active mechanism, with the receiver cell actively adapting to the signal from the sender cell. We then demonstrate how the degree of active coupling is controlled by key morphological parameters, such as junction length and the degree and orientation of mechanical polarization. Strikingly, force propagation is amplified perpendicularly to the axis of mechano-structural polarization, similar to the Poisson effect in passive material. Finally we verify that our findings in these cell doublets can be generalized to larger cell clusters. Overall, we show that active cellular responses to incoming forces can maintain signal strength and leads to the emergence of an apparent elastic behaviour that allows signals to be propagated over large distances, as in an elastic material, rather than be dissipated, as in a viscous material.

## Results

### The intercellular junction decreases the mechano-structural polarization

The most important feature of epithelial tissue is strong cell-cell adhesion, which makes the epithelial monolayer a coherent sheet that can effectively separate different compartments, like the outside and inside of a body or organ. Therefore, we first characterised how the presence of an intercellular junction influences cellular organization and force generation. To this end, we compared single cells (“singlets”) and cell pairs (“doublets”) grown on identical micropatterns (Fig. 1 A). The H-pattern is known to be able to accomodate both singlets and doublets, which in both cases form an hour-glass shape.

**Figure 1:**
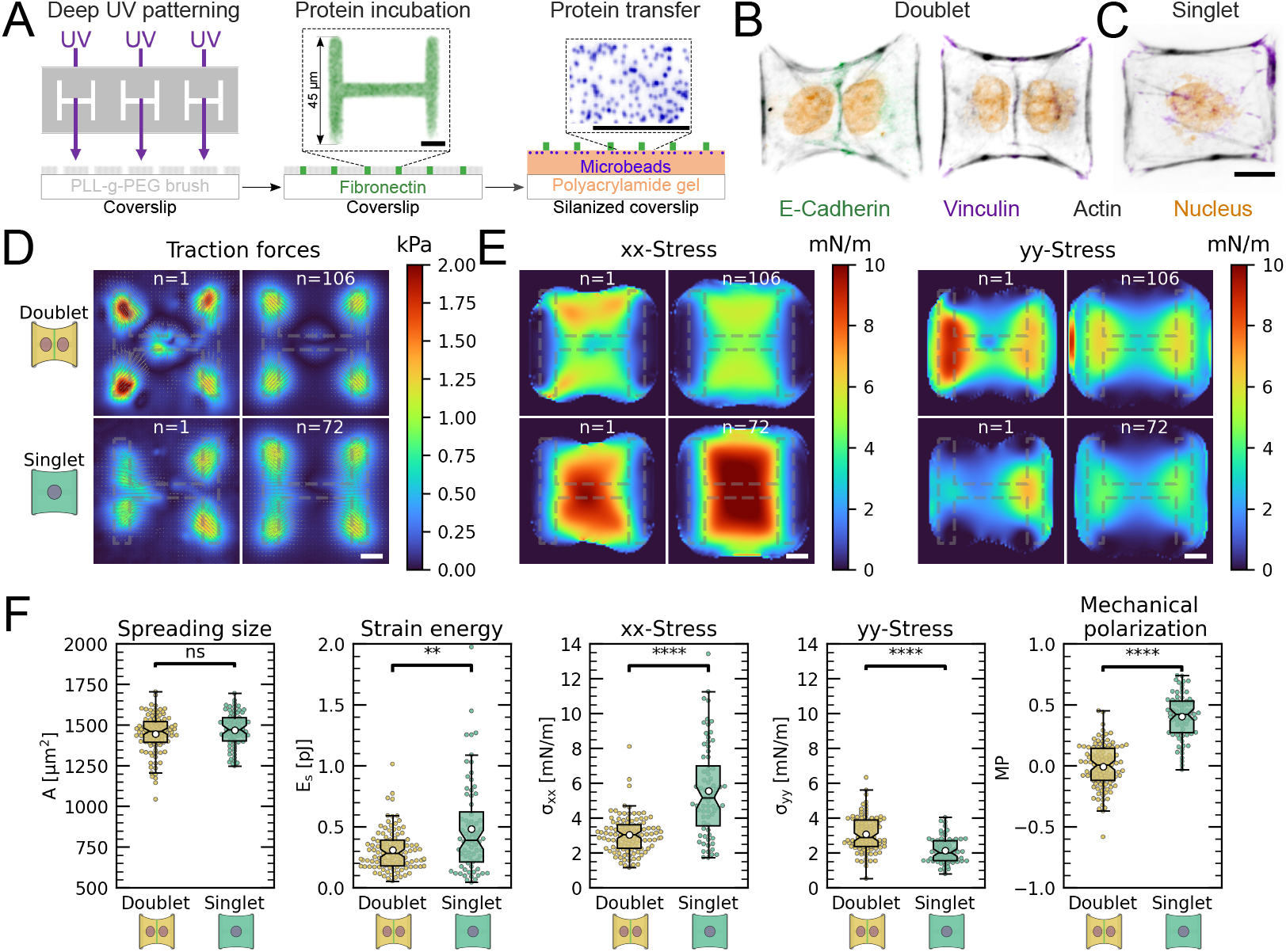
The cell-cell junction leads to a decrease in mechanical polarization. **A** Cartoon of the micropatterning process on soft substrates, allowing to control cell shape and to measure forces at the same time by embedding fluorescent microbeads into the gel and measuring their displacement. The middle panel shows the used pattern geometry, an H with dimensions of 45 μmx45 μm. **B, C** Immunostaining of opto-MDCK cells plated on H-patterns and incubated for 24 h before fixing. Actin is shown in black, E-Cadherin in green, Vinculin in violet and the nucleus in orange. **B** The left and right images show a representative example of a doublet **C** A representative example of a singlet. **D** Traction stress and force maps of doublets (top) and singlets (bottom) with a representative example on the left and an average on the right. **E** Cell stress maps calculated by applying a monolayer stress microscopy algorithm to the traction stress maps, with a representative example on the left and an average on the right. **F** From left to right, boxplots of: Spreading size, measured within the boundary defined by the stress fibers. Strain energy, calculated by summing up the squared scalar product of traction force and displacement field divided by two. xx-Stress and yy-stress calculated by averaging the stress maps obtained with monolayer stress microscopy. Degree of polarization, defined as the difference of the average xx- and yy-stress normalized by their sum. Doublets are shown in yellow and singlets are shown in green. The figure shows data from n=106 doublets from N=10 samples and n=72 singlets from N=12 samples. All scale bars are 10 μm long.

We found that when plated on H-shaped micropatterns, singlets formed prominent stress fibers around the cell contour (peripheral stress fibers), as well as some smaller internal stress fibers which resulted from the spreading process (Fig. 1 C and Fig. S1). Vertical stress fibers at the edge of the patterns, along the vertical bars of the H, were straight and strongly coupled to the substrate (adherent stress fibers), while peripheral stress fibers located above the non-adhesive regions of the micropattern, in between the vertical bars of the H, were curved due to the inward pull of the cell cortex (free stress fibers). Focal adhesions were primarily located in the corners of the pattern, although some were present on the middle bar of the H-pattern, which is required for the cells to spread over the whole pattern. A similar pattern of organization was observed in doublets, with the addition of a prominent cell-cell junction in the center of the H-pattern, parallel to the lateral bars of the H (Fig. 1 B), consistent with previous work [27].

By quantifying cell-generated forces using traction force microscopy (TFM), we found that the magnitude of traction forces is surprisingly similar between singlets and doublets (Fig. 1 D). When we quantified the overall contractility by calculating the strain energy stored in the substrate, we found that it is even slightly higher for singlets than for doublets, despite spreading over the same surface area (Fig. 1 F). This is likely because singlets have to spread a smaller volume over the same surface as doublets, leading to higher tension. Moreover they do not have to accomodate any cell-cell junction and therefore could be coupled better to the substrate [27].

Next we calculated stresses born by the cells using monolayer stress microscopy (MSM), which converts the TFM-data into an estimate for intracellular stress (Fig. 1 E). In doublets, the normal stresses in x- and y-direction (*σ*_**xx**_ and *σ*_**yy**_) were comparable, whereas in singlets *σ*_**xx**_ was much larger than *σ*_**yy**_ (Fig. 1 F). To quantitatively compare the cellular stress distribution of these systems, we computed the mechanical polarization as (*σ*_**xx**_ – *σ*_**yy**_)/(*σ*_**xx**_ + *σ*_**yy**_). With this quantification, a system polarized vertically has a polarization of −1, 0 reflects an unpolarized system and 1 a horizontally polarized system. Doublets were unpolarized (average degree of polarization of 0), whereas singlets were horizontally polarized with an average degree of polarization of almost 0.5. Next, we measured the polarization of the actin structures with a homemade algorithm using the structure tensor (see methods section for details). We found the same trend and a strong correlation between mechanical and structural polarization, meaning that the stress fibers in singlets are largely organized horizontally, whereas in doublets they are directed more towards the center (Fig. S2). Our results suggest that intercellular junctions may act as a barrier preventing the horizontal organization of stress fibers that exist in singlets, thus strongly altering the mechanical polarization of the system.

### The presence of an intercellular junction leads to a redistribution of tension from free to adherent peripheral stress fiber

An inherent limitation of TFM is that it only quantifies tension transmitted to the substrate while forces internally balanced are not detected. Although this is partially remedied by MSM, which estimates an internal stress distribution based on the TFM-results, this method lacks spatial resolution to take into account the precise shape of the cell. In order to address this important aspect, we therefore turn to contour models that focus on the role of the peripheral stress fibers (Fig. 2 A).

**Figure 2:**
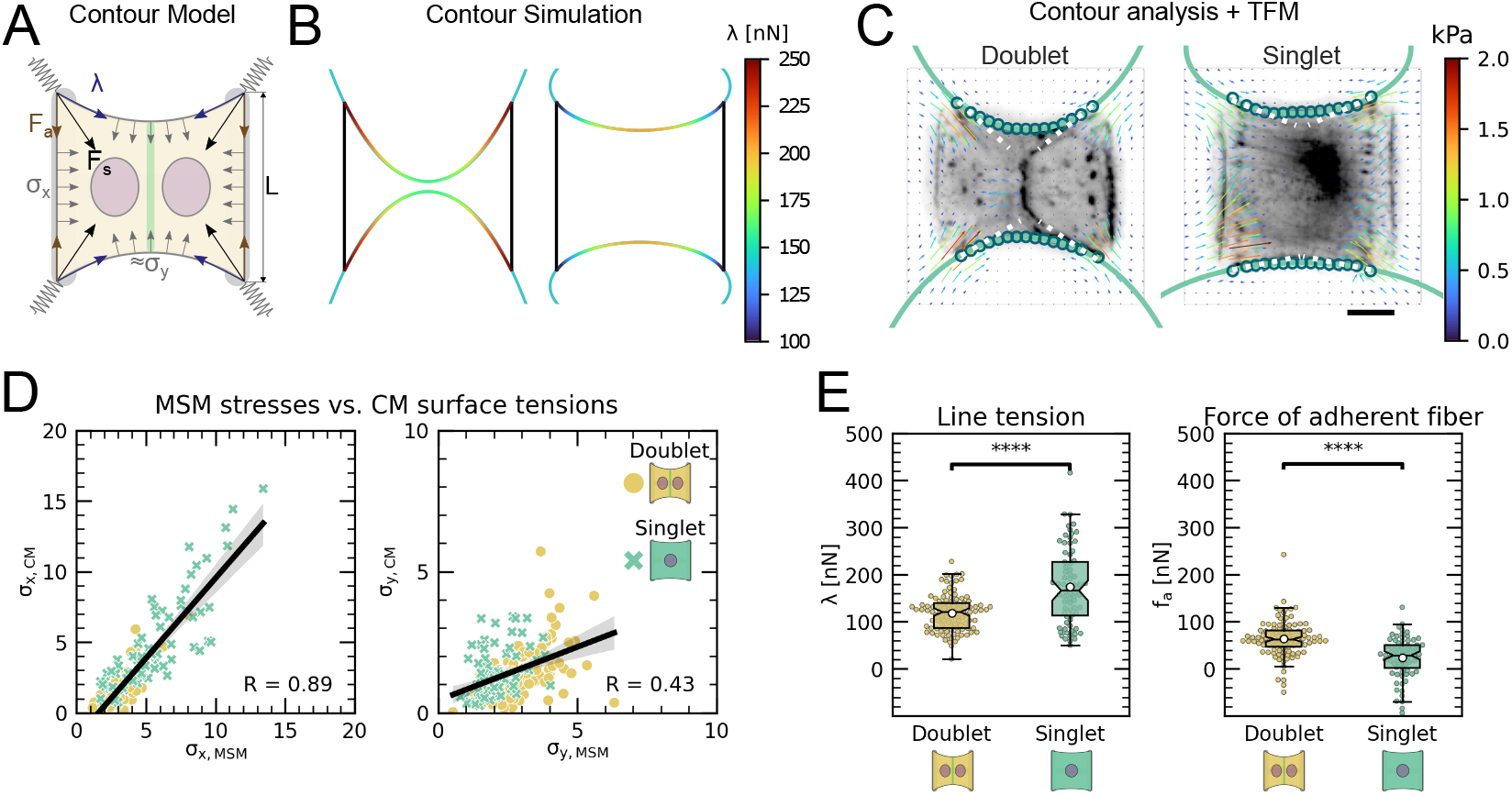
The cell-cell junction leads to a redistribution of tension from free to adherent peripheral stress fiber. **A** Cartoon of the contour model used to analyze the shape of the doublets and singlets. **B** FEM simulation of the contour with *σ_y_ > σ_x_* left and *σ*_x_ > *σ*_y_ right. **C** Actin images of doublets (left) and singlets (right) with traction stresses (arrows), tracking of the free fiber (blue circles), elliptical contour fitted to the fiber tracks (green line) and tangents to the contour at adhesion point (white dashed line). The scale bar is 10 μm long. **D** Correlation plot of MSM stresses and CM surface tensions. MSM stresses were calculated by averaging the stress maps obtained with monolayer stress microscopy and the surface tensions were obtained by the contour model analysis, where *σ*_x_ was measured on the TFM maps by summing up the x-traction stresses in a window around the center of the vertical fiber and *σ*_y_ was determined by fitting the resulting ellipse to the tracking data of the free fiber. Doublets are shown as yellow dots and singlets are shown as green crosses. The black line shows the linear regression of the data and the shaded area shows the 95% confidence interval for this regression. The R-value shown corresponds to the pearson correlation coefficient. **E** Boxplots of line tension *λ* (left) and force of adherent fiber F_a_ (right) as defined in panel A. Both values were calculated by first calculating the force in each corner by summing up all forces in a radius of 12 μm around the peak value and then projecting the resulting force onto the tangent of the contour for the line tension and onto the y-axis for the force of adherent fiber. Doublets are shown in yellow and singlets are shown in green. The figure shows data from n=106 doublets from N=10 samples and n=72 singlets from N=12 samples. All scale bars are 10 μm long.

We previously showed that the curvature of a free stress fiber results from a balance between an isotropic surface tension pulling the stress fibers towards the cell center and a line tension acting along the fibers, tending to straighten them [28, 29]. The radius of curvature is then given by the ratio of the line to the surface tension. As the line tension can be calculated from the TFM data and the radius of curvature can be measured, the surface tension can be inferred. One key assumptions of our previous work was that cellular tension is isotropic. As we showed that single cells are mechanically polarized (Fig. 1 F), we generalized our circular arc model to anisotropic systems [30], allowing to compute surface tensions in the x- and y-direction by measuring the surface tension in x-direction on the TFM maps and then fitting the surface tension in y-direction until the resulting ellipse fits to the fiber (Fig. 2 A and supplementary theory). Application of this approach to experimental data allowed to compute surface tensions for both singlets and doublets (Fig. 2 B, C).

The combination of MSM and contour analysis allow thorough characterization of cell mechanics. MSM describes the bulk mechanics of cells assuming they are linear elastic and ignoring peripheral stress fibers, while contour analysis mostly characterises the peripheral stress fibers and assumes they are linearly elastic only under tension, like a rubber band.

Analysis using both of these methods demonstrates that stress fibers in singlets are subjected to a larger stress along the x-direction than in doublets and conversely that stress fibers in doublets are subjected to higher stresses in the y-direction than singlets (Fig. 2 D). Consistent with this, singlets possessed a significantly larger line tension in their free stress fibers than doublets. In contrast, the force exerted by adherent stress fibers displayed the opposite behaviour: it was higher in doublets than in singlets (Fig. 2 E). These two forces were computed by integrating the traction stresses in each corner, correcting for the contribution of the surface tension along the adherent fiber and then projecting these forces onto the stress fibers (see supplemental theory for details). These results are consistent with the MSM analysis Fig. 1. Indeed, *σ*_**xx**_ (which corresponds roughly to the free stress fiber, since it is approximately parallel to the x-axis) is higher in singlets and *σ*_**yy**_ (which corresponds roughly to the adherent stress fiber, since it is parallel to the y-axis) is higher in doublets (Fig. 1 E). We conclude that the presence of cell-cell junction leads to a redistribution of tension from the free to adherent peripheral stress fibers.

### Force increase through local activation of RhoA in one cell leads to active force increase in neighboring cell in doublets

In order to study signal propagation, it is important to generate a well-defined input whose propagation can be followed in space and time. Although this is a notoriously difficult issue in cellular force generation, a new tool was recently established which allows just that, namely non-neuronal optogenetics. In order to switch on cell contractility in a controlled manner, we activated RhoA, a major regulator of cell contractility, with an optogenetic actuator that relocalises a RhoGEF domain to the membrane in response to 488 nm light (Fig. 3 A [31]).

**Figure 3:**
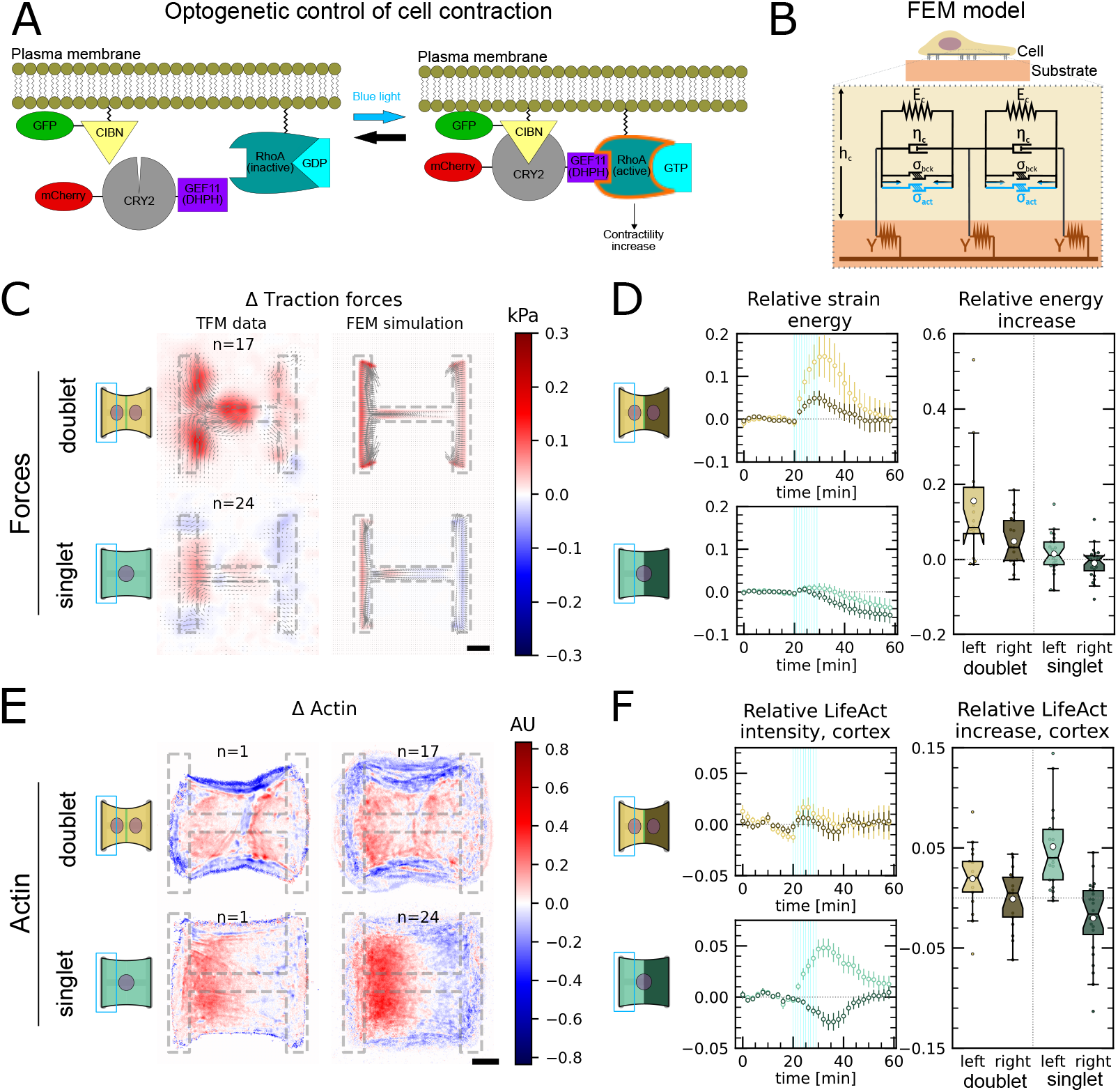
Local activation of RhoA leads to stable force increase in both the activated and the non-activated cell in doublets, but destabilizes force homeostasis in singlets. **A** Cartoon of the optogenetic CIBN/CRY2 construction used to locally activate RhoA. **B** Cartoon of the FEM model used to explain optogenetic experiments. **C** Difference of average traction force maps after and before photoactivation of cell doublets (top) and singlets (bottom). Maps on the left show the TFM data and maps on the right show the result of the FEM simulations with an active response of the right cell. **D** Relative strain energies of doublets (top) and singlets (bottom) with local photoactivation, divided in left half (bright) and right half (dark). One frame per minute was acquired for 60 minutes and cells were photoactivated with one pulse per minute for 10 minutes between minute 20 and minute 30. Strain energy curves were normalized by first substracting the individual baseline energies (average of the first 20 minutes) and then dividing by the average baseline energy of all cell doublets/singlets in the corresponding datasets. Data is shown as circles with the mean ±s.e.m. Boxplots on the right show the value of the relative strain energy curves 2 minutes after photoactivation, i.e. at minute 32. **E** Difference of actin images after and before photoactivation of doublets (top) and singlets (bottom), with an example on the left and the average on the bottom. All scale bars are 10 μm long. **F** LifeAct intensity measurement inside the cells over time (left) of left half (bright) vs right half (dark) of doublets (top) and singlets (bottom) after local photoactivation. Boxplots on the right show the relative actin intensity value after 2 minutes after photoactivation of activated vs non-activated half. The figure shows data from n=17 doublets from N=2 samples and n=17 singlets from N=6 samples. All scale bars are 10 μm long.

As previous work has shown that this tool allows localized activation of signalling within single cells ([32]) and we used it to activate the left half of singlets and doublets to determine how the localized stress created by activation propagated to the other side of the system. To make sure we do not accidentally activate the right cell, we estimated how much stray light the right cell receives, photoactivated the whole cell with this power density and found no measurable contraction of the doublet (Fig. S3, Movie S7). Surprisingly, the stress propagation differed markedly between singlets and doublets. In doublets, traction forces increased both in the activated and the non-activated region. In the singlets, on the other hand, traction forces increased slightly and very locally in the activated region, but decreased in the non-activated region (Fig. 3 C-D, Movie S4, Movie S5). We conclude that in contrast to singlets, doublets can establish stable contraction patterns under half-activation of contractility.

We hypothesised that this surprising behaviour may originate from differences in the re-organization of contractile elements within the cytoskeleton in singlets and doublets. Therefore, we imaged the behavior of the actin cytoskeleton during the light stimulation by comparing the fluorescence intensity distribution of the F-actin reporter LifeAct before and during stimulation. In doublets, LifeAct fluorescence increases slightly inside and decreases slightly outside of the doublet. The decrease outside of the doublet is mostly due to fiber movement. When we measure the LifeAct intensity following its movement, the intensity remains mostly constant (Fig. S2 C). In contrast, in singlets, LifeAct fluorescence redistributed from the unstimulated side to the stimulated side, both inside of the cell as well as on the periphery (Fig. 3 E-F, Fig. S2 C).

To determine if the behavior of the doublets could arise from a passive response of the non-activated region, we developed a finite element (FE) model to predict stress propagation (Fig. 3 B). Based on previous work characterising cell rheology [33, 34, 35, 36], our model consists of a network of Kelvin-Voigt elements that are each connected to an elastic substrate. Each Kelvin-Voigt element also possesses an active element, which describes the contractility of myosin motors that can be increased to simulate optogenetic activation of contractility. In order to fix the parameters of the model, we performed an experiment where we photoactivated the whole singlet/doublet (see supplementary theory and Movies S1-S3 for details). We used this model to predict the spatiotemporal evolution of traction stress in the system. Comparison of the FE results to the experimental data shows that the behavior of the non-activated region cannot be reproduced with a purely passive reaction (Fig. S4 A-B). Therefore, we hypothesised that active coupling takes place, perhaps due to mechanotransductory signalling pathways. To test this idea, we introduced an active coupling element into the FEM-model between the left and the right half. We then used this coupling term as a fitting parameter to qualitatively reproduce the experimental traction maps. Again, singlets differed from doublets. Coupling in doublets was positive, meaning the right half contracts in response to the contraction of the left half; whereas it was negative in singlets, meaning the right half relaxes in response to the contraction of the left half.

Together, these data indicate that cells in the doublet are actively coupled, with the unstimulated cell responding to the contraction of the stimulated cell by actively contracting, in agreement with previous qualitative reports [37, 38]. Strikingly, traction force generated by doublets shows a homeostatic response to this transient increase of RhoA activity. Indeed, once activation is stopped, the traction force generated on the pattern returns to its initial level. In singlets on the other hand, transient and local RhoA activation has a destabilizing effect. The local increase in traction stress and the local accumulation of F-actin in the photoactivated region is compensated with a decrease in stress and F-actin in the nonactivated region. Furthermore, rather than displaying a homeostatic behaviour, the traction stress keeps decreasing even after the activation is stopped. We hypothesize, that this may occur because the actin structures acutely fluidize in response to the local stress increase, as previously reported [39, 40]. Since there is no junction and thus no diffusion barrier in singlets, the imbalance in stress induced by optogenetic activation may lead to a flow of Factin from the non-activated to the activated region, consistent with our observations (Fig. 3 E-F). As a qualitative test, we exchanged the Kelvin-Voigt elements in our model of the cell body for Maxwell elements after photoactivation. This led to a behaviour consistent with our observations (Fig. S5 C-D).

Overall, our data show that the cytoskeleton possesses active coupling and that the degree of coupling depends on the presence of an intercellular junction. The intercellular junction allows efficient propagation of stress across the whole micropattern, probably due to mechanotransductory pathways and by impeding fluidization.

### Strong active coupling is present in the actin cortex of doublets

Having shown that the unstimulated cell in doublets reacts actively to the contraction of the stimulated cell, we sought to quantify the strength of this active response. To this end, we sought to quantitatively reproduce the distribution of cell stresses obtained by MSM in photoactivated doublets using our FEM-model (Fig. 4 A-C). To simulate optogenetic activation, we increased the level of contractility of the activated left hand side of the doublet compared to the baseline found in unstimulated conditions. Then to simulate coupling, we tuned the degree of contractility on the unstimulated right hand side of the doublet. The ratio of contractility of the right half to the left half corresponds to the degree of active coupling between the cells in the doublet. An active coupling of 0 means no contraction of the right half, 1 indicates a contraction of the right half of the same magnitude as the left and −1 means relaxation of the right half with same magnitude as the increase on the left. To allow comparison of experiments to simulations, we normalize the stress increase of the right cell by the total stress increase (Fig. 4 C). For each experiment, we determined the degree of coupling that best reproduced the experimental cellular stress distribution in the x- and y-directions (Fig. 4 B).

**Figure 4:**
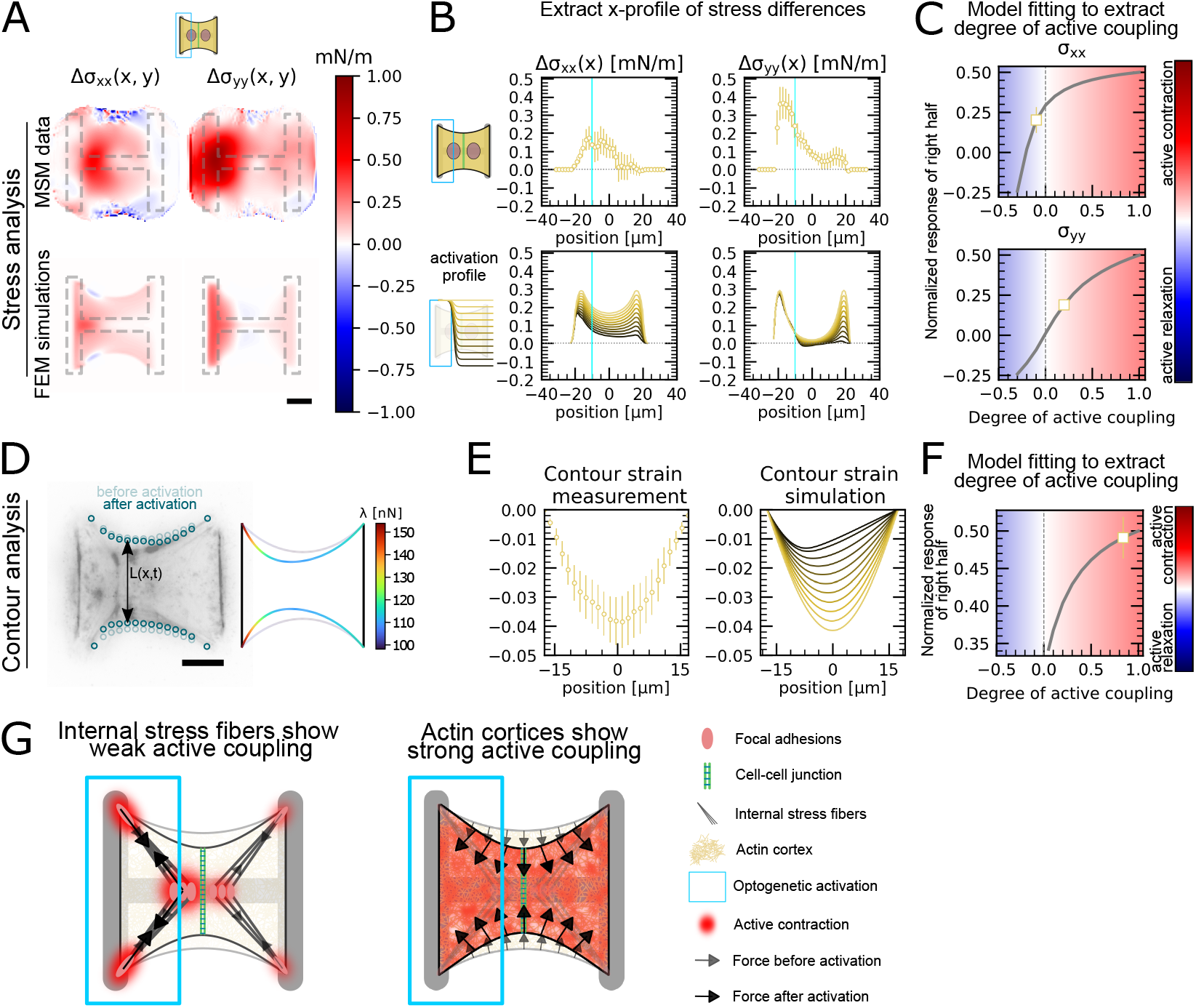
Stress and contour modelling show strong active coupling of actin cortices in doublets. **A** Difference of average cell stress maps after and before photoactivation of cell doublets, calculated with MSM (top) and simulated in an FEM model (bottom) Stress in x direction is shown on the left and stress in y direction is shown on the right. **B** Average over the y-axis of the maps in A. Data is shown as circles with the mean ±s.e.m. In the simulation, the right half of the cell was progressively activated to obtain the family of curves shown in the bottom. **C** Response of the right half (normalized by the total response), obtained from the model (grey line), as a function of the degree of active coupling. The experimental MSM value is placed on the curve to extract the degree of active response of the right cell in the experiment. **D** Contour analysis of the free stress fiber. In the experiment, the distance between the free fibers as a function of x is measured, as shown in the image on the left. An example for a contour model simulations is shown in the right. **E** The contour strain after photoactivation is calculated from the distance measurements shown in D, by dividing the distance between the free stress fibers for each point in x-direction after and before photoactivation. Similarly to the FEM simulation, in the contour simulation, the right half of the contour is progressively activated to obtain the curve family shown in the right plot. **F** Response of the right half (normalized by the total response), obtained from the model (grey line), as a function of the degree of active coupling. The experimental strain value is placed on the curve to extract the degree of active response of the right cell in the experiment. **G** A cartoon showing our interpretation of the results shown in panel A to F. The traction force analysis only measures forces that are transmitted to the substrate, which are dominated by the activity of the stress fibers. The contour of the free fiber is determined by the activity of the actin cortex and the free stress fiber. Thus, the strong active coupling in the contour suggests strong active coupling of the cortices and the comparatively weak active coupling of the forces suggests a weak active coupling of the stress fibers. The figure shows data from n=17 doublets from N=2 samples. All scale bars are 10 μm long.

Interestingly, this analysis showed different coupling behaviours in the x- and y-directions. We found positive active coupling in the y-direction (0.2), but negative coupling in the x-direction (−0.05) (yellow square, Fig. 4 C). This may be because all forces in y-direction are balanced between the cell and the substrate, but not across the junction. This signifies that each cell can contract independently from one another in this direction. In contrast, the forces in the x-direction must always be balanced by interaction between the cells across the junction, similar to a “tug of war”.

To test our hypothesis of independent contraction in the y-direction, we measured the distance between the free stress fibers along the x-axis (Fig. 4 D) to get a readout for cortical tensions not transmitted to the substrate. The ratio of the inter-stress fiber distance during and before photoactivation defines a contour strain along the x-direction (Fig. 4 E). We compared experimental contour strain to the contour strain in simulations, in which we again progressively activated the right half of the contour (Fig. 4 D-E), and repeated the same analysis as in Fig. 4 C. We found a degree of coupling of 0.8, indicating a global active contraction of the unstimulated cell (Fig. 4 F). This is consistent with the active positive coupling measured in the y-direction using MSM Fig. 4 C. Overall both TFM and surface tension analyses showed active coupling between the two regions. However, active coupling was weaker in TFM measurements, perhaps because the cortices of the two cells are more strongly actively coupled than the stress fibers.

In conclusion, traction forces, as measured by TFM, show weaker active coupling between activated and non-activated region than cortical tensions, as inferred by measurement of contour strain. The traction forces are dominated by the activity of the stress fibers, both internal and on the periphery, because most forces are found in the corners of the doublet. The only area where the cortex can transmit forces to the substrate is along the vertical fiber in horizontal direction. If this force were substantial, it should point much more horizontally and be much more constant, without the strong hotspots in the corners. The contour of the free fiber on the other hand is determined by the activity of the actin cortex and the free stress fiber. Thus, contour analysis suggests strong active coupling of the cortices and the comparatively weaker active coupling observed in cellular stress distributions may occur because internal stress fibers are coupled to the substrate and transmit little stress across the cell junction (Fig. 4 G).

### Mechanical stresses transmit most efficiently perpendicularly to the axis of mechanical and structural polarization in doublets

Our data indicated that active coupling of contractions in the y-direction is much higher in doublets than in the x-direction. We hypothesized that active coupling may be modulated by mechanical and structural polarization of the cells. To test this, we sought out to vary structural and mechanical polarization of doublets by changing the aspect ratio of the underlying micropatterns from 1to2, 1to1, to 2to1 (y to x ratio) while maintaining a constant spreading area. Mechanical polarization and structural polarization was quantified as previously. We found that structural and mechanical polarization are tightly correlated and vary greatly in between the three different aspect ratios (Fig. 5 A-C). For example, on micropatterns with 1to2 aspect ratio, both stress fibers and force patterns were oriented horizontally whereas on 2to1 they were oriented vertically.

**Figure 5:**
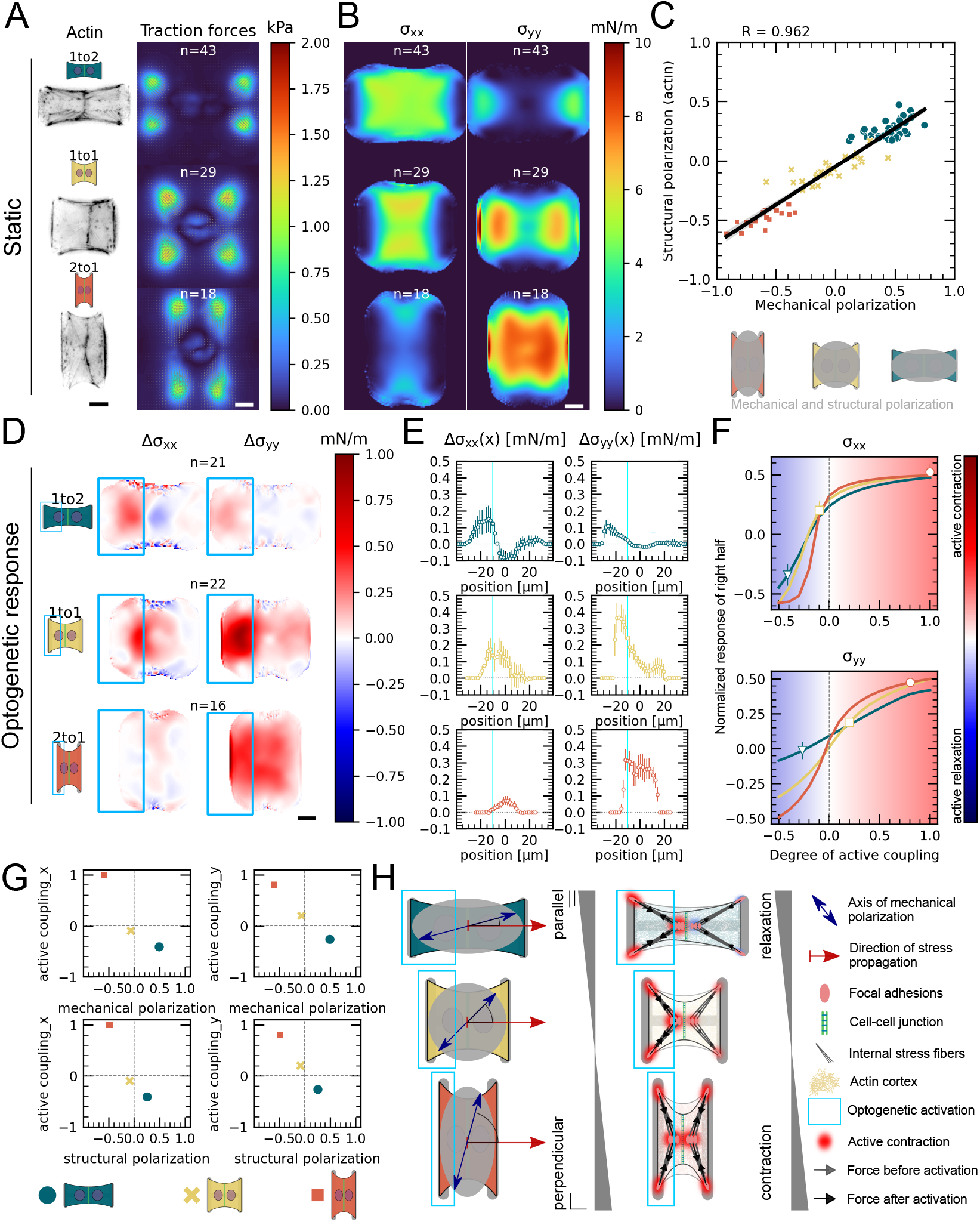
Mechanical stresses transmit most efficiently perpendicularly to the axis of mechanical and structural polarization in doublets. **A** Actin images (left) and average traction stress and force maps (right) of cell doublets on H-patterns with different aspect ratios (1to2, 1to1 and 2to1). **B** Average cell stress maps calculated by applying a monolayer stress microscopy algorithm to the traction stress maps. **C** Correlation plot of mechanical and structural polarization. The black line shows the linear regression of the data and the shaded are shows the 95% confidence interval for this regression. The R-value shown corresponds to the pearson correlation coefficient. **D** Stress maps of the difference of xx-stress (left) and yy-stress (right) before and after photoactivation. **E** Average over the y-axis of the maps in D. Data is shown as circles with the mean ±s.e.m. **F** Response of the right half (normalized by the total response), obtained from the model (grey line), as a function of the degree of active coupling. The experimental MSM value is placed on the curve to extract the degree of active response of the right cell in the experiment. All scale bars are 10 μm long. **G** The degree of active coupling plotted against the average mechanical and structural polarization **H** A cartoon showing our interpretation of the data shown in panel A to F. The relative response of the right cell in response to the activation of the left cell varies strongly in the different aspect ratios. In the 1to2 doublet, where polarization and transmission direction are aligned, the right cell relaxes, whereas in the 2to1 doublet, where the polarization axis is perpendicular to the transmission direction, the right cell contracts almost as strongly as the left cell. The figure shows data from n=43 1to2 doublets from N=6 samples, n=29 1to1 doublets from N=2 samples and n=18 2to1 doublets from N=3 samples. For the analysis of the optogenetic data, doublets with unstable stress behavior before photoactivation were excluded. All scale bars are 10 μm long.

Next, we examined the link between structural polarization and stress transmission. For each aspect ratio, we repeated the local activation experiments (Fig. 4, Fig. 5 D-F, Movie S6). These optogenetically induced stresses transmit from the sender cell to the receiver cell, i.e. from left to right. We observed markedly different behaviour depending on aspect ratio. In 1to2 doublets, cells are polarized mechanically and structurally along the direction of stress transmission and, after activation of left hand cell, the right cell reacts by relaxing.

In contrast, in 2to1 doublets, cells are polarized mechanically and structurally perpendicular to the direction of stress transmission and activation of the left hand cell leads to contraction of the right hand cell. We then computed the degree of active coupling as previously and found that the degree of active coupling increased with increasing mechanical and structural polarization (Fig. 5 D-G).

We then investigated whether a similar effect could be observed for cortical tensions and performed the contour analysis as in Fig. 4. Here we saw, in agreement with Fig. 4 E, that the contour deformation is very symmetrical in both the 1to1 and the 2to1 doublets, but much less in the 1to2 doublets, where the degree of active coupling is lower. The quantification of the degree of active coupling here is lower for the 2to1 than for the 1to1, but the uncertainty of this quantification is quite high because the contour strain is small, so this is likely due to the noise in the strain measurements (Fig. S5). Altogether, we conclude that mechanical stresses transmit most efficiently perpendicularly to the axis of mechanical and structural polarization in doublets (Fig. 5 G).

### Mechanical stresses transmit most efficiently perpendicularly to the axis of mechanical and structural polarization in small cell clusters

Next we investigated whether this conclusion is generalizable to larger systems, such as small monolayers. To this end, we confined about 10-20 cells on 150 μmx40 μm rectangular micropatterns. We again performed TFM and MSM experiments as well as live imaging of Factin and quantified the mechanical and structural polarization for micropatterns with aspect ratios of 1:4. We observed prominent actin cables at the periphery of the small monolayers with less marked stress fibers internally. In these conditions, the tissue is mechanically and structurally polarized along the long axis of the pattern (Fig. 6 A-C, Fig. S6). We then characterised the efficiency of stress propagation parallel and perpendicular to the axis of tissue polarization. To this end, we photoactivated either the top half or the left half of the tissues. In our experiments, we observed again an increase of traction forces and cell stress both in the activated and in the non-activated region. We computed the degree of active coupling in the same way as for doublets using our FEM-model and found that active coupling is higher, when the direction of stress propagation is perpendicular to the axis of mechanical and structural polarization of the tissue. Additionally, we measured the distance *d* over which the stress attenuates to 20% of its maximum, and found that *d* is, on average, three-fold larger when the direction of stress propagation is perpendicular to the axis of polarization (Fig. 6 D-F). We conclude the correlation between mechano-structural polarization and active coupling observed in doublets is also present in larger groups of cells. In summary, active coupling and its correlation with mechanical and structural polarization seems to be typical for epithelia, independent of size (Fig. 6 G).

**Figure 6:**
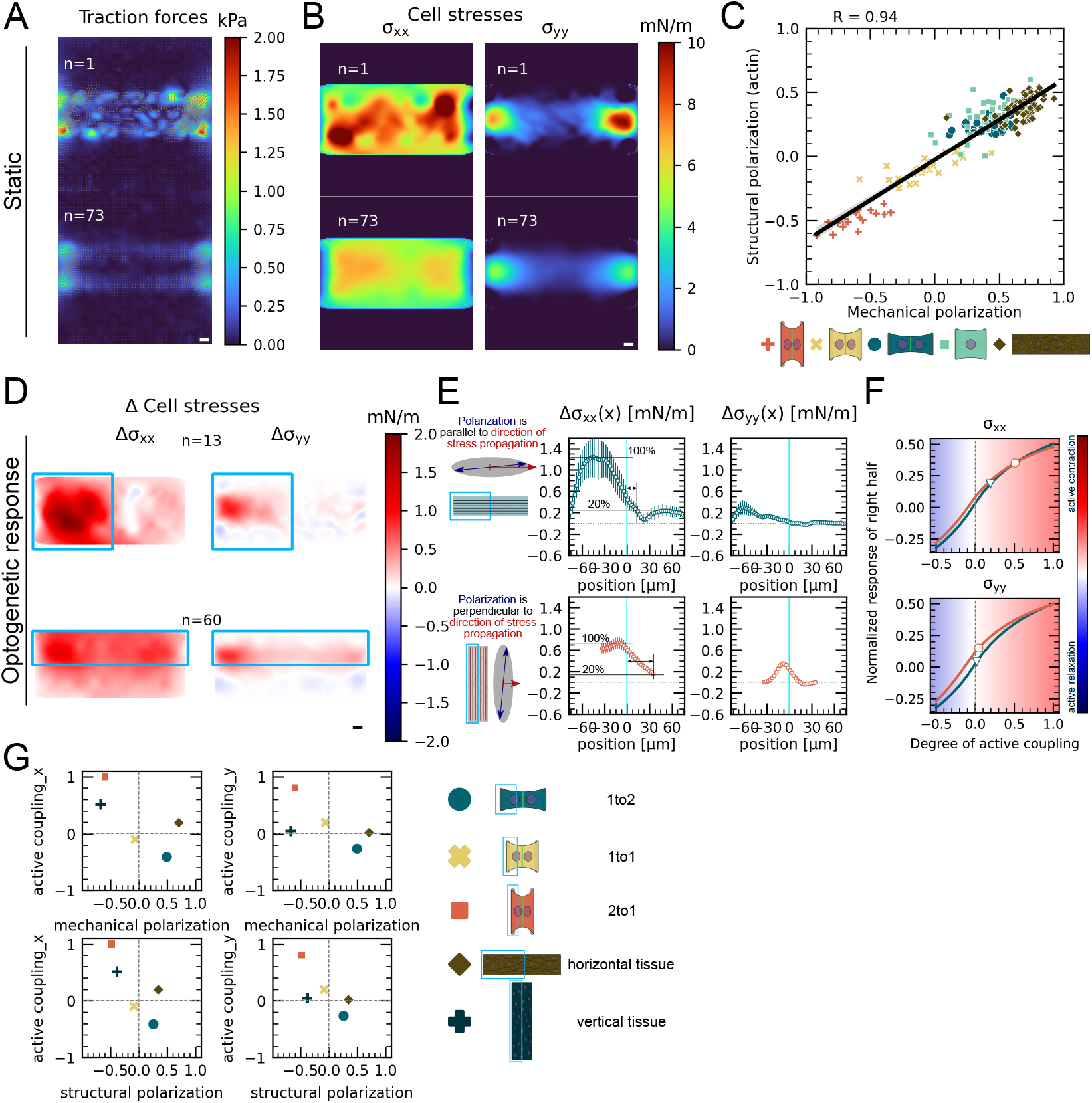
Mechanical stresses transmit most efficiently perpendicularly to the axis of mechanical and structural polarization in small monolayers. **A** Representative (top) and average (bottom) maps of traction forces and stresses of a small monolayer on rectangular micropattern. **B** Representative example and average cell stress maps calculated by applying a monolayer stress microscopy algorithm to the traction stress maps. **C** Correlation plot of mechanical and structural polarization across all conditions. The black line shows the linear regression of the data and the shaded are shows the 95% confidence interval for this regression. The R-value shown corresponds to the pearson correlation coefficient. **D** Stress maps of the difference of xx-stress (left) and yy-stress (right) before and after photoactivation. **E** Average over the y-axis of the maps in D. Data is shown as circles with the mean ±s.e.m. **F** Response of the right half (normalized by the total response), obtained from the model (grey line), as a function of the degree of active coupling. The experimental MSM value is placed on the curve to extract the degree of active response of the right cell in the experiment. **G** The degree of active coupling plotted against the average mechanical and structural polarization. All scale bars are 10 μm long. The figure shows data from n=13 tissues from N=2 samples photoactivated on the left and from n=60 tissues from N=3 samples photoactivated on the top. All scale bars are 10 μm long.

## Discussion

Intercellular forces play a major role in regulating and coordinating tissue morphogenesis. Recent work has shown that mechanical forces participate in long-range signalling, propagating over large distances at which they can be received and interpreted by other cells [**?**]. However, we have little quantitative insight of how cell-generated forces propagate across intercellular junctions or which cellular structures modulate propagation. This is because direct measurement of intercellular forces and cell internal stresses within embryos or tissues is very challenging and most of our knowledge of the distribution of these forces is inferred from theoretical models [41, 42, 43, 44]. By combining quantitative measurements of cellular stresses and cell shape with optogenetic control of contractility and mathematical modelling, here we showed that force signal propagation within cellular assemblies is an active process whose amplification mechanism is controlled by the mechanostructural polarization of the system.

Our results revealed the presence of active coupling between cells. This was demonstrated before by Liu et al. [45], but a thorough quantification of this active coupling has been lacking. Photoactivation of one cell in our doublet leads to contraction, sending a force signal. The receiver cell reacts to this signal with an active contractile response. We quantified the response of the receiver cell by comparing experimental traction force and cell stress data with an FEM model. We found, that a purely passive reaction of the receiver cell cannot account for the data and therefore concluded that the receiver cell reacts actively. This active coupling mechanism increases the spatial range a mechanical signal can travel to about 1 or 2 cell lengths according to our data. Furthermore, analysis of the cell shape showed very high symmetry of shape deformation despite the asymmetrical photoactivation. This shape deformation is dominated by the activity of the actin cortex and comparison of this measurement with a mathematical contour model lead us to conclude that the active coupling of the cortices is stronger than that of stress fibers. However, this is probably strongly influenced by tissue and cell mechanical properties and by geometry and mechanical properties of the substrate. Additionally, we tested only transient signals. Maintaining signal strength over longer periods of time could also lead to farther transmission of force signals. Compared to chemical signals, these mechanical signals can travel very fast: Indeed, with our temporal resolution of 1 frame per minute, no delay between receiver and sender cell was apparent. In contrast, when we carried out the same activation protocol on a single cell that had the same area as the doublets, the non-activated region displayed acute fluidization of the actin structure. Thus, in the absence of an intercellular junction, localized contraction leads to actin flow instead of the stress buildup observed in doublets. Therefore, cellularization of the tissue may allow compartmentalization of stress and efficient transmission of stress, allowing the tissue to act as an elastic material rather than a viscous fluid.

Several subcellular features determined the efficiency of active coupling. Indeed, our experiments revealed that intercellular coupling strongly depends on the anisotropy of F-actin organization and force distribution. We found, that the magnitude of contraction of the receiving cells relative to the sender cells depends on the direction and magnitude of its mechanostructural polarization. If the tissue’s or doublet’s polarization axis is perpendicular to the axis between sender and receiver cells, the receiver cells react more strongly and the signal travels farther. However, determining the exact contribution of subcellular structures remains challenging because the cell forms a highly coupled system comprising dynamic mechanotransduction feedback loops. Future work will be necessary to determine the molecular mechanisms detecting the mechanical signal, transducing it, and amplifying it. In particular, it will be interesting to investigate how the active contraction of the receiver cell depends on its own mechanostructural polarization and that of the sender cell. Currently, the nature of the stimulus detected by the receiver cell is unclear. We note that mechanics and biochemistry are closely coupled, because strain can change biochemistry by changing concentrations and spatial localization, and stress on single molecules can open cryptic binding sites or increase dissociation constants.

Finally our study of epithelial monolayers show that the supracellular organization of actin is a major regulator of force propagation within tissues. Forces are transmitted more efficiently in a direction perpendicular to the axis of actin polarity also in small monolayers. These results give rise to several interesting conclusions. First, recent studies have proposed that groups of cells can behave as a “supracellular unit”, which share many of the characteristics of the individual cells that it consists of [46, 47, 48]. Some emerging mesoscale phenomena, such as collective gradient sensing, might be explained by common principles, such as supracellular polarity and supracellular force transmission [49, 50, 51, 52, 53]. Our findings complement those results, as we show that the correlation between mechano-structural polarization and force signal transmission distance holds true across scales. Second, at a much larger scale, we speculate that propagation through active coupling may have important implications in developmental processes, such as convergent extension in the xenopus mesoderm. In these tissues, cells are planary polarized in a direction perpendicular to the extension of the tissue, and the convergence and extension of the tissue is driven by directed contraction and migration of the cells [54]. Our results suggest that preferential transmission of active contraction perpendicular to the polarization axis of the cells could amplify this mechanism and contribute to the robustness of the process.

## Supplementary figure captions

**Fig. S1**

Immunostaining of opto-MDCK cells plated on H-patterns and incubated for 24 h before fixing. Actin, E-Cadherin, Vinculin and the nucleus is shown. The top and middle row show a representative example of a doublet and the bottom row shows a representative example of a singlet. Scale bar is 10 μm long.

**Fig. S2**

**A** LifeAct images of some example opto-MDCK doublets (left) and singlets (right) plated on H-patterns. **B** Correlation plot of mechanical and structural polarization. The black line shows the linear regression of the data and the shaded are shows the 95% confidence interval for this regression. The R-value shown corresponds to the pearson correlation coefficient. Scale bar is 10 μm long.**C** LifeAct intensity measurement on periphery of cells over time (left) of left half (bright) vs right half (dark) of doublets (top) and singlets (bottom) after local photoactivation. Boxplots on the right show the relative actin intensity value after 2 minutes after photoactivation of activated vs non-activated half

**Fig. S3**

**A** Fluorescence image of a homogeneously coated coverslip, illuminated on a rectangular zone with a DMD and the horizontal intensity profile. The background was subtracted from the image before making the measurements. The peak intensity drops to 6% over a distance of 10 μm, so all local activation routines were placed 10 μm away from the junction. **B** Left: A cartoon describing the experiment. First, the whole doublet is activated at low light intensity (0.054 mW mm^-2^) and then only the left cell is activated with higher intensity (0.9 mW mm^-2^). The intensity of the first activation routine corresponds to the intensity right at the junction from the second activation routine. Right: Curves show the strain energy stored in the substrate under the left cell (left) and under the right cell (right). The strain energy curves were normalized by the strain energy level before photoactivation. Red curves show the average and grey curves show the individual doublets.

**Fig. S4**

**A** Difference of average traction force maps after and before photoactivation of cell doublets (top) and singlets (bottom). Maps on the left show the TFM data and maps on the right show the result of the FEM simulations without any active reaction of the right cell. **B** Relative strain energies of doublets (top) and singlets (bottom) with local photoactivation, divided in left half (bright) and right half (dark). One frame per minute was acquired for 60 minutes and cells were photoactivated with one pulse per minute for 10 minutes between minute 20 and minute 30. Strain energy curves were normalized by first substracting the individual baseline energies (average of the first 20 minutes) and then dividing by the average baseline energy of all cell doublets/singlets in the corresponding datasets. Data is shown as circles with the mean ±s.e.m. and simulated curves are shown as solid lines. **C** A cartoon showing the basic elements of the FEM simulation. Acute fluidization is modeled as a switch from Kelvin-Voigt to Maxwell elements. **D** Relative strain energies of singlets with local photoactivation, ddivided in left half (bright) and right half (dark). One frame per minute was acquired for 60 minutes and cells were photoactivated with one pulse per minute for 10 minutes between minute 20 and minute 30. Strain energy curves were normalized by first substracting the individual baseline energies (average of the first 20 minutes) and then dividing by the average baseline energy of all cell doublets/singlets in the corresponding datasets. Data is shown as circles with the mean ±s.e.m and the result of an FEM simulation is shown as a solid line.

**Fig. S5**

**A** Contour analysis of the free stress fiber. In the experiment, the distance between the free fibers as a function of x is measured, as shown in the image on the left. **B** The contour strain after photoactivation is calculated from the distance measurements shown in A, by dividing the distance between the free stress fibers for each point in x-direction after and before photoactivation. **C** Response of the right half (normalized by the total response), obtained from the model (solid lines), as a function of the degree of active coupling. The experimental strain value is placed on the curve to extract the degree of active response of the right cell in the experiment.

**Fig. S6**

Phalloidin stainings of actin structures of small tissues. Scale bar is 10 μm long.

**Movie S1**

Actin + traction forces (left) and relative strain energy (right) over time of a globally photoactivated doublet.

**Movie S2**

9 examples of globally photoactivated doublets. Actin is shown in black, traction forces are overlaid as colored arrows, the tracked contour in blue circles, the tangents in white dashed lines and the fitted ellipse in green

**Movie S3**

9 examples of globally photoactivated singlets. Actin is shown in black, traction forces are overlaid as colored arrows, the tracked contour in blue circles, the tangents in white dashed lines and the fitted ellipse in green

**Movie S4**

Actin + traction forces (left), traction force map (center) and relative strain energy divided in left (blue) and right (orange) half (right) over time of a locally photoactivated doublet.

**Movie S5**

Actin + traction forces (left), traction force map (center) and relative strain energy divided in left (blue) and right (orange) half (right) over time of a locally photoactivated singlet.

**Movie S6**

Average traction force maps (top) and relative strain energy divided in left (bright) and right (dark) half (right) over time of locally photoactivated 1to2 (blue), 1to1 (yellow) and 2to1 doublets (red).

**Movie S7**

Cry2 distribution with photoactivation of left cell in a doublet with increasing power densities. 1st pulse: 0.18mW mm^-2^, 2nd pulse: 0.9mW mm^-2^, 3rd pulse: 1.8mW mm^-2^, 4th pulse: 3.6 mW mm^-2^, 5th pulse: 9mW mm^-2^, 6th pulse: l8mWmm^-2^

## Materials & Methods

### Cell Culture

Opto-MDCK and opto-MDCK LifeAct cells have been kindly provided by Manasi Kelkar and Guillaume Charras. Both cell lines were cultured at 37 °C and in 5% CO_2_ atmosphere in DMEM (Life Technologies) medium containing 10% heat-inactivated FBS (Life Technologies) and 1% penicillin/streptomycin (Sigma-Aldrich). Between 20.000 and 50.000 cells were plated on the micropatterned hydrogels. After 1 h, cells were checked for their adhesion to the hydrogels. In case of excessive amount of cells the sample was rinsed with fresh medium to wash off the non-adhered cells. Cells were let spread on patterns for 16 h to 28 h, so that on average most doublets have started as single cells and divided on the pattern to form a doublet.

### Cell fixing and immunostaining

First, cells were fixed for 10 min with 4% PFA diluted in PBS. Next, the cell membrane was permeabilized with 0.5% Triton X-100 for 5 min. Cells were then washed twice with TBS and blocked at room temperature for 1 h with a blocking buffer solution containing TBS, 1% bovine serum albumin (BSA, Sigma-Aldrich) and 50mM Glycine (Sigma-Aldrich). Then, cells were incubated for 2 h in a dilution of primary antibodies with blocking buffer. For E-Cadherin stainings a 1:200 dilution of DECMA-1 (ThermoFisher 14-3249-82) was used and for Vinculin stainings a 1:400 dilution of hVIN-1 (Sigma-Aldrich V9131) was used. Cells were then washed three times with TBS for 10 min each. Then cells were incubated in a dilution of secondary antibodies, Alexa 555-conjugated phalloidin and DAPI in blocking buffer. For E-Cadherin stainings a 1:1000 dilution of Alexa 647-conjugated anti-rat (Sigma-Aldrich SAB4600186) was used, for Vinculin stainings a 1:1000 dilution of Alexa 647-conjugated anti-mouse (ThermoFisher A-21235) and a 1:1000 dilution for phalloidin and DAPI. Fixed cells were then mounted with Mowiol 4-88 (Polysciences, Inc.) onto glass slides and kept at 4 °C until imaging.

### Preparation of micropatterned polyacrylamide gels

Patterned PAA hydrogels were prepared according to the glass method described previously in [55]. In short, 32mm coverslips were first plasma cleaned for 60 s and then incubated with a drop of PLL-PEG 0.1 mg mL^-1^ in HEPES 10mM, ph 7.4 for 30 min at room temperature. Then, coverslips were rinsed with a squirt bottle of MilliQ water and carefully dried with a nitrogen gun. The coverslips were then placed on a quartz photomask (Toppan) on a 10 μL drop of MilliQ water. Excess water was removed by placing a kimwipe on the coverslips, a flat surface on top (e.g. the lid of a petridish) and then pressing gently. The coverslips on the photomask were then exposed to deep-UV for 5 min. After recovery from the photomasks, the coverslips are incubated with 20 μgmL^-1^ fibronectin (Sigma-Aldrich) and 20 μg mL^-1^ Alexa488-conjugated fibrinogen (Invitrogen) in 100mM Sodium Bicarbonate buffer for 30min at room temperature. To prepare the gels, a 47 μL drop of 20 kPa mix of polyacrylamide and bis-acrylamide (Sigma-Aldrich) was prepared (see [56] for the proportions). To perform Traction Force Microscopy, carboxylate-modified polystyrene fluorescent microbeads (Invitrogen F-8807) were added to the polyacrylamide premix and sonicated for 3 min to break bead aggregates. A second coverslip of the same size is then placed on top, after previous silanization with a solution of 5mL 100% ethanol, 18.5 μL Bind Silane (GE Healthcare Life Science) and 161 μL 10% acetic acid (Sigma-Aldrich) for 5 min. During the polymerization process, the hydrogel adheres to the silanized coverslip and fibronectin proteins are trapped within the polyacrylamide mesh. The silanized coverslip is finally detached by wetting it with MilliQ water, letting the gel rehydrate for 5 min and lifting it up with a scalpel. Hydrogels were stored in 100 mM Sodium Bicarbonate buffer at 4 °C for maximum 2 days before cell seeding.

### Imaging and optogenetic photoactivation

All experiments were conducted 16 h to 28 h after seeding the cells on the sample. Then the cells were observed on an inverted Nikon Ti-E2 microscope with an Orca Flash 4.0 sCMOS camera (Hamamatsu), a temperature control system set at 37°C, a humidifier and a CO_2_ controller. For the opto-experiments on cell doublets and singlets a Nikon 60x oil objective was used and for the opto-experiments on tissues a Nikon 40x air objective was used. The E-cadherin and vinculin staining images were taken with an Eclipse Ti inverted confocal microscope (Nikon France Instruments, Champigny sur Marne, France), equipped with sCMOS prime camera (Photometrics), a 60× objective, and a CSU X1 spinning disk (Yokogawa, Roper Scientific, Lisses, France). MetaMorph software was used for controlling the microscope (Universal Imaging Corporation, Roper Scientific, Lisses, France). Unless otherwise stated, all photoactivations were done with 1 pulse per min for 10 min and each pulse had a duration of 200 ms, a power density of 0.9 mW mm^-2^ and a wavelength of 470 nm. The power density was measured with a power meter right after the objective by shining light on a surface of a given size and dividing the measured power by this size.

### Traction Force Microscopy and Monolayer Stress Microscopy

Force measurements were performed using a method described previously [57]. In short, fluorescent beads were embedded in a polyacrylamide substrate with 20 kPa rigidity and images of those beads were taken before, during and after photoactivation. At the end of the experiment, cells were removed with 2.5% Trypsin and an unstressed reference image of the beads was taken. The displacement field analysis was done using a homemade algorithm based on the combination of particle image velocimetry and single-particle tracking. After correcting for experimental drift, bead images were divided into smaller subimages of 13.8 μm width. The displacement between corresponding bead sub-images was obtained by crosscorrelation. After shifting the stressed sub-images to correct for this displacements, the window size is divided by 2 and new displacement values are determined by cross-correlations on the smaller sub-images. This procedure is repeated twice. On the final sub-images, singleparticle tracking was performed: this ensures that the displacement measurement has the best possible spatial resolution at a given bead density. Erroneous vectors were detected by calculating the vector difference of each vector with the surrounding vectors. If the vector magnitude was higher than 2.5 μm or the vector difference higher than 1 μm, the vector was discarded and replaced by the mean value of the neighbouring vectors. Only the first frame of each movie was compared to the unstressed reference image. All subsequent frames were compared to their predecessor. This leads to more precise measurements because the displacements are much smaller. From the bead displacement measurements a displacement field was then interpolated on a regular grid with 1.3 μm spacing. Cellular traction forces were calculated using Fourier transform traction cytometry with zero-order regularization [58] [59], under the assumption that the substrate is a linear elastic half-space and considering only displacement and stress tangential to the substrate. To calculate the strain energy stored in the substrate, the scalar product of the stress and displacement vector fields was integrated over the surface of the whole cell. The algorithm was implemented in MATLAB. Cell internal stresses were calculated from the traction stress with the code from Bauer et al. [60]. To do this calculation, the cell is assumed to behave like a thin, elastic sheet that is attached to a substrate and then contracts. Equilibrium shape is reached, when the active stress that leads to the contraction is balanced by the elastic stress that builds up within the sheet and in the substrate. The resulting stress is the sum of the active and the passive stress in the elastic sheet and is independent of its elastic modulus.

### Fiber Tracking

A semi-automatic procedure was used to detect and track the actin fibers at the cell contour over time. First the operator clicks on the endpoints of each fiber on the first image of a time lapse. The adherent fibers are very static and straight, so, in this case, we just draw a straight line between the two end points. The free fibers are curved and move over time. To follow the shape of a given fiber over time, we used a custom script: on each image, parallel line profiles are drawn at regular intervals in between the two defined endpoints, in a direction perpendicular to the overall fiber direction; each profile is analyzed to detect the point where it intersects the fiber, using intensity variation as criterion. The line linking these points describes the actin fiber position at each time point. In order to filter out badly detected points, the consistency of the resulting positions is analyzed over both time and space. Temporal filtering consists of first a median filter over 5 time points and the removal of outliers. Within a moving time window of 10 time points, positions distant from the average value by more than 2 times the standard deviation are deleted. Spatial filtering includes also removal of outliers, defined as being distant from the spatial average position by more than 3 times the standard deviation. Then the angle of lines joining adjacent points are computed at each position and badly tracked points are excluded by ensuring that these angles stay below 15°. Finally, we use this tracking data to create a stack of masks for each cell which accurately describes the complete contour of the cell. The algorithm was implemented in MATLAB.

### Actin polarization analysis

To measure the average polarization of the internal actin network, we analyze the orientation of the internal actin network using the structure tensor formalism. For each pixel with intensity *I*(*x,y*), the structure tensor *J* is calculated over a Gaussian local neighborhood *w*(*x, y*) with a waist of 3 pixels, according to equation (1).

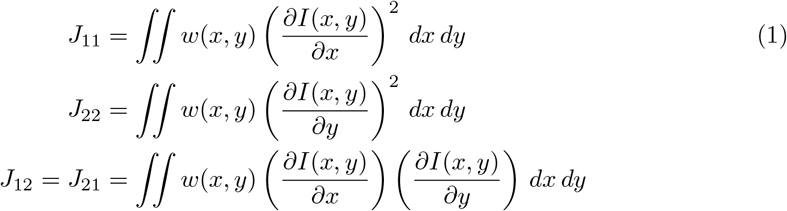

The orientation angle *θ* on this local neighborhood corresponds to the direction of the main eigenvector of the structure tensor and is obtained by equation (2).

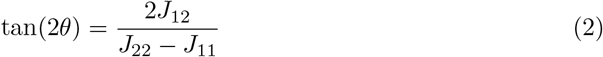

This angle is only meaningful if the image shows oriented structures in this neighborhood. This confidence can be estimated from the coherency, which quantifies the degree of anisotropy and is calculated from the structure tensor according to equation (3). Values with a coherency value under 0.4 were excluded before averaging the orientation angles over the cell to obtain the mean direction of the actin network. The degree of polarization is then obtained according to (4) The algorithm was implemented in MATLAB.

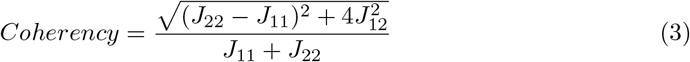

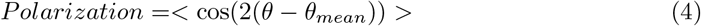

### Actin Intensity Measurement

To measure the actin intensity in the left and the right half of the doublet/singlet, we first segment the cells using the masks obtained from the fiber tracking. We reduce its size a little bit to exclude the external stress fibers from the measurement. We then divide the doublet/singlet vertically in two halves and sum up all the intensity values within the region of interest, yielding one intensity value per frame and per half. This intensity over time is then normalized by the intensity value of the average over the first 20 frames before photoactivation.

### Statistical analysis and boxplots

All boxplots show the inner quartile range as boxes and the whiskers extend to 1.5 times the inner quartile range. The notches show the 95 % confidence interval for the median and the white dot shows the sample mean. The Mann-Whitney-Wilcoxon U test was used to test for differences between singlets and doublets, with ns: p > 0.05, *: p < 0.05, **: p < 0.01,***: p < 0.001 and ****: p < 0.0001.

### Data exclusion for optogenetic experiments

Many of the cells showed an unstable baseline energy level, which made it difficult to judge the impact of the optogenetic activation. Thus, we quantified the baseline stability of each cell by applying a linear regression to the relative strain energy curve before photoactivation and excluded all cells with a slope larger in absolute value than a threshold value. For figure 3, this process excluded 16 globally activated doublets, 7 globally activated singlets, 12 locally activated doublets and 17 locally activated singlets. For figure 5 D to F, this process excluded 22 1to2 doublets, 7 1to1 doublets and 2 2to1 doublets.

## Author contributions

A.R. performed experiments from figures 1 to 5, a large part of the data analysis and writing the manuscript under the supervision of M.B., T.B., G.C. and U.S.S.. D.W. developed the contour model, the finite element model and developed the algorithms for the circle and ellipse fitting under the supervision of U.S.S.. V.M. performed the experiments and data analysis from figure 6 under the supervision of M.B.. M.K. developed the optogenetic cell line expressing LifeAct under the supervision of G.Ch.. I.W. developed the TFM algorithm, the algorithm for measuring the orientation of the actin network and the algorithm for tracking the actin fibers. P.M. designed the photomasks used to make the micropatterns and, together with J.R., provided technical assistance and maintenance for the technical facilities. All authors contributed with editing the manuscript and providing fruitful discussions and feedback.

## Acknowledgements

M.B. acknowledges financial support from the French Agence Nationale de la Recherche (ANR) MechanoSwitch project, grant ANR-17-CE30–0032-01. U.S.S. acknowledges funding through a joint ANR-DFG grant, project number SCHW 834/2-1. T.B. acknowledges fundings through CNRS grants (Actions Interdisciplinaires 2017, DEFI Instrumentation aux limites 2017, Tremplin@INP 2021, PEPS CNRS-INSIS 2021). G.Ca. acknowledges financial support from the ANR SupraWaves project, grant ANR-19-CE13-0028. This work was supported by the Center of Excellence of Multifunctional Architectured Materials “CEMAM” (n° AN-10-LABX-44-01).

We would like to thank Luis Vigetti and Isabelle Tardieux for providing access to their spinning disc confocal microscope that was used to take the immunofluorescence images. We would like to thank Simon de Beco, Laurent Blanchoin and the whole MicroTiss team for useful discussions and feedback.

## 1 Two-dimensional continuum modelling of cellular contractility

This mesoscopic model approximates the cell as an elastic continuum. The general constitutive relation can be written as

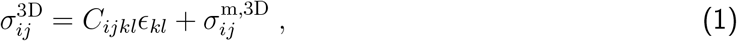

with total stress tensor 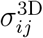, stiffness tensor *C_iykl_*, strain tensor *ϵ_kl_* and motor stress tensor 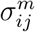. Further, the force balance equation

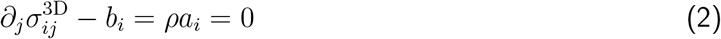

is used to calculate the deformation of the cell, where *b_i_* is the external body force acting on the cell. For cells or tissues we always assume the inertial term to vanish.

### 1.1 Thin-layer approximation

We next assume that we make is that the effective thickness of the cell is much smaller than the overall extent of the cell *h_c_* ≪ *L_c_*. Thus, variations along the z-direction are assumed to be small and it is sufficient to consider a thickness-averaged stress tensor given by

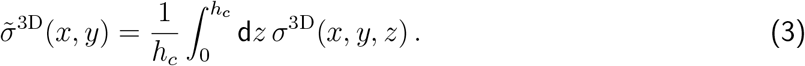

Averaging the force balance equation leads to a two-dimensional force balance equation in which the thickness-averaged body force is now acting as a traction

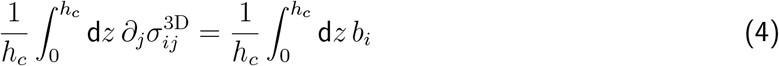

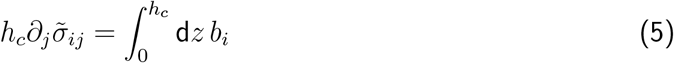

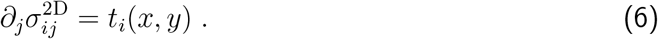

The cell thickness is the conversion factor between three-dimensional and two-dimensional quantities, *q*^2D^ = *q*^3D^*h_c_*.

### 1.2 Plane stress

Under plane stress assumption we set *σ_zz_* = *σ_xz_* = *σ_zx_* = *σ_yz_* = *σ_zy_* = 0 and further neglect out-of-plane strain *ϵ_zz_*. Hooke’s law under plane stress conditions can be written in Voigt notation as

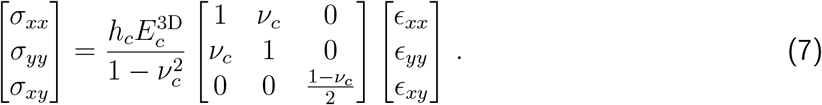

Together with the general version of Hooke’s law

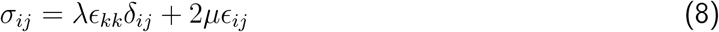

we determine the 2D Lamé parameter as

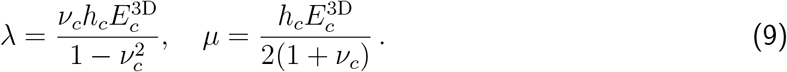

### 1.3 Active Kelvin-Voigt model

The constitutive relation of an active Kelvin-Voigt model in index notation is given by

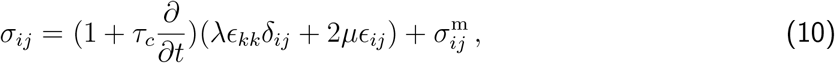

with stress tensor *σ_ij_*, strain tensor *ϵ_ij_* and the 2D Lamé coefficients as defined in equation (9). The material relaxation time is defined as *τ_c_* = *η_c_*/*E_c_* with *η_c_* denoting the cell viscosity. The linearized strain tensor is defined as

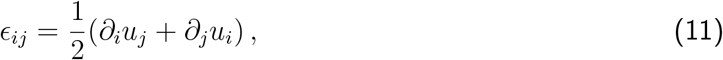

where *U_j_* is the *j*^th^ component of the displacement field vector **u**(**x**). The overall active contraction is described by the anisotropic motor stress tensor 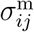 which is split into

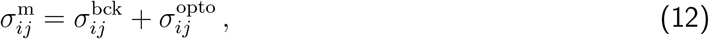

i.e. a time-independent background stress to account for the cellular energy baseline level and a time dependent photo-activation stress tensor describing the stress increase during photo activation (PA). Based on experimental observations and verification with the MSM analysis of the TFM data, the anisotropy of the cytoskeleton enters the stress tensor for the background stress through the mechanical polarization which is defined as

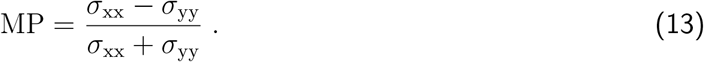

This leads to

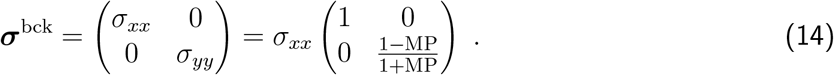

Upon photo-activation we assume a time dependent stress contribution given by

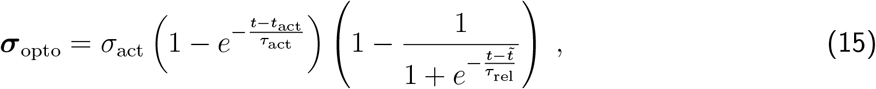

which is a combination of an increasing saturating exponential and a sigmoidal shaped decrease (Fig. 1AS).

### 1.4 Cell - substrate coupling

The cell substrate coupling is described by equation (6) where the traction is formulated as

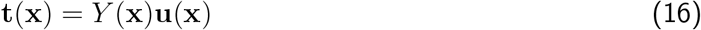

which yields

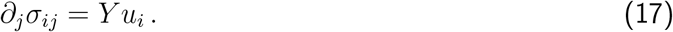

*Y* denotes the position-dependent spring stiffness density. Combining equation (10) and one can show that the interplay of cellular and substrate elasticity defines a natural length scale

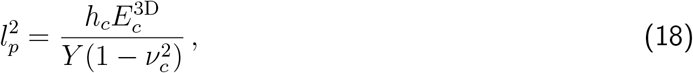

known as the force-localization length which describes how far a point force is transmitted in the elastically coupled isotropic material. The stiffness of the substrate can be estimated via

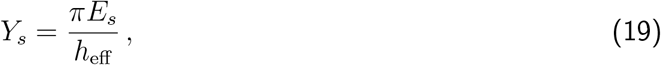

in which the effective substrate height is given by an interpolation formula

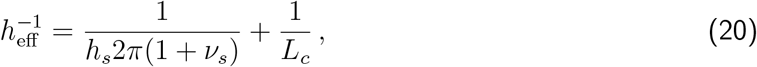

where *h_s_* and *L_c_* denote the substrate height and cell layer size, respectively. *E_s_* and *ν_s_* are the Young’s modulus and Poisson’s ratio of the substrate. To adapt our theory as close as possible to the traction force computation of the experiments we assume that the substrate is infinitely thick and therefore we have *h*_eff_ ≈ *L_c_*. Further, we have for the traction forces at the cell-substrate interface

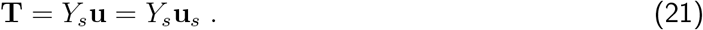

The elastic energy stored in the substrate is calculated via

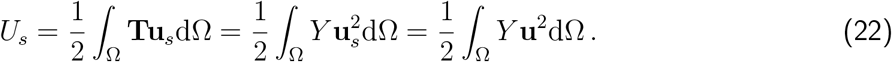

### 1.5 Parametrization

Although in principle it is possible to use a downhill-simplex method to find the set of parameters which minimizes the theoretically computed substrate energy against the experimentally measured curve, we nevertheless decide to fix some of the parameters to avoid overfitting. All fixed parameters are listed in Tab. 1. While the substrate parameters are known, we fix the parameters for Young’s modulus and viscosity of the cell to typically reported values from the literature [1, 2, 3, 4, 5]. The fixed substrate parameters yield a spring stiffness density of *Y_s_* = 1.257 × 10^9^Nm^−3^ and a force-localization length of *l_p_* = 3.25 μm. Simulations with these parameters lead to a very good agreement of theoretically computed and experimentally measured stress and traction maps. Although the traction and stress maps as computed with the FEM-model show all characteristic features of the experimental maps, latter seem to be blurred. The blurring can be traced back to the TFM and MSM analysis methods.

**Table 1:** Fixed parameters for the two-dimensional finite element simulation.

3 Supplement Figures and Tables
| Fixed parameter | Value |
| --- | --- |
| Substrate |  |
| Young's modulus of the substrate $E_s$ | 20 kPa |
| Poisson's ratio of the substrate $\nu_s$ | 0.5 |
| Thickness of the substrate $h_s$ | 50 $\mu\text{m}$ |
| Cell |  |
| Young's modulus of cell $E_c$ | 10 kPa |
| Viscosity of the cell $\eta_c$ | 100 kPa s |
| Thickness of the cell $h_c$ | 1 $\mu\text{m}$ |
| Poisson's ratio of the cell $\nu_c$ | 0.5 |
| Length of the cell $L_c$ | 50 $\mu\text{m}$ |

### 1.6 Finite Element Simulation

We solve the combination of equations (10) and (17) for the displacement vector **u** of the cell by means of a finite element simulation using the open source software package FEniCS [6]. This approach has been used in several other works [1, 2, 7, 8, 3, 4, 9]. The full problem statement is given by: Find the displacement field vector **u**(**x**)with initial conditions **u**_0_ = **u**(**x**, 0) = 0 such that together with 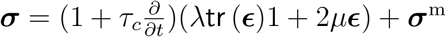

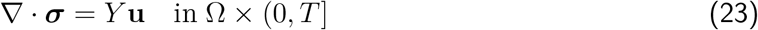

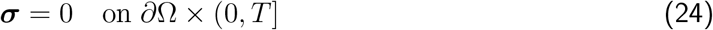

Therefore, we derive the weak form of equation (17) by multiplying with a vector-valued test function 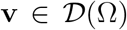 over the simulation domain Ω. Multiplying equation (23) with the test function and integrating over the whole simulation domain leads to

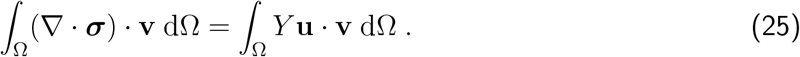

The left hand side can be integrated using integration by parts i.e. using the following identity

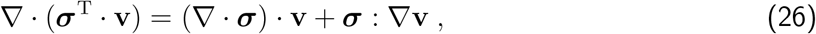

where we use the standard notation for the inner product between tensors (double contraction) and ∇**v** = *∂_i_* (*v_j_* **e**_*j*_) ⊗ **e***_i_* being the vector gradient. Equation (25) can be simplified to

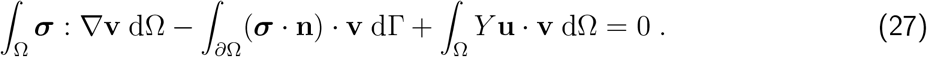

***σ*** · **n** is the traction vector at the boundary ⌈ = *∂*Ω which is set to zero in case of stress free boundaries. We further use that ***σ*** is symmetric and thus, the double contraction with the antisymmetric part 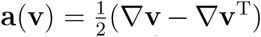 of ∇**v** is zero i.e. ***σ***: **a**(**v**) = 0. This allows us to replace ∇**v** by its symmetric part 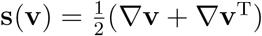 and leads to the final weak form statement

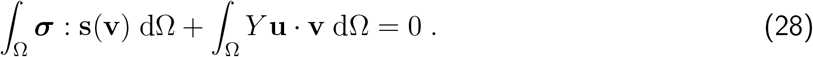

Since we are aiming at solving for the displacment vector **u** we have to express all terms in the constitutive relation in terms of **u**

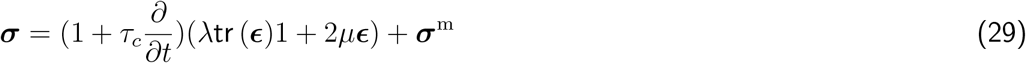

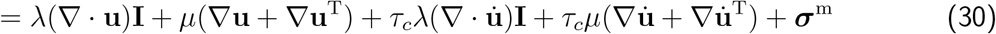

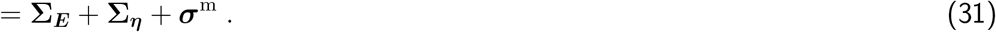

For the time derivatives we use a backward Euler discretization scheme which is numerically stable even for larger time steps. We set

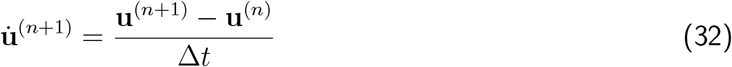

and since we are dealing with linear equations the discretization scheme translates directly to

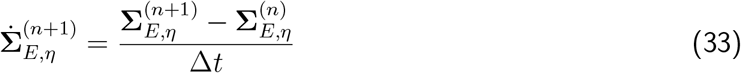

which enables us to define

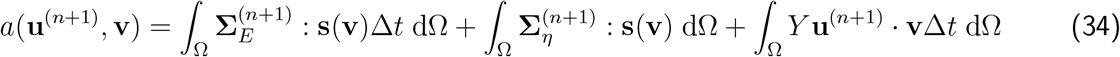

and

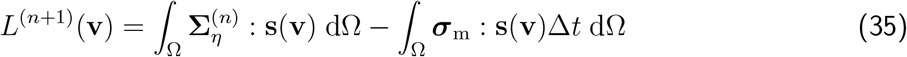

after inserting the time discretized version of equation (31) in equation (28). Our initial problem statement now reduces to solving

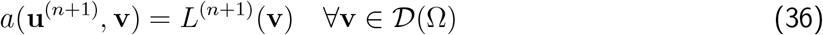

Equation (36) can be directly handed to the FE solver.

### 1.7 Modelling procedure for the 2D finite element simulation

To obtain the final theoretical result by means of our finite element simulation (Fig.4C), several steps were necessary. We used the open source meshing software GMSH [10] to create a finite element mesh as depicted in Fig. 1BS. Then we fixed all known parameters in order to match the experimental setup to our simulations. All fixed parameters are gathered in Tab. 1 and were fixed throughout the simulations. Next we mathematically defined the pattern geometry of the H-pattern which determines the portion of the simulation domain on which the cell is assumed to establish a connection to the elastic foundation (Fig.1CS)

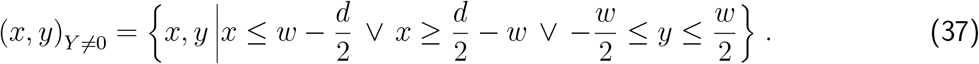

We first determined the active background stress by fitting the baseline of strain energy curve (Eq. 22) to the given experimental substrate strain energy. In a second step we fitted the temporal evolution of the strain energy by optimizing the free parameters *σ*_act_, *τ*_act_ and 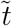 in Eq. 15.

The obtained parameters for all fitted conditions are summarized in Tab. 2. The fit results of the doublet and singlet strain energy curves can be see in Fig. 4D.

**Table 2:** Fit parameter as obtained by the two-dimensional finite element simulation.

| Fit parameter | Singlet | Doublet |
| --- | --- | --- |
| Baseline |  |  |
| Background stress component $\sigma_{xx}^{\text{bck}}$ | 6.59 kPa | 5.73 kPa |
| Background stress component $\sigma_{yy}^{\text{bck}}$ | 2.78 kPa | 5.73 kPa |
| Full opto-stimulation |  |  |
| Active stress $\sigma_{\text{act}}$ | 0.287 kPa | 0.618 kPa |
| Activation time scale $\tau_{\text{act}}$ | 133 s | 227 s |
| Relaxation time scale $\tau_{\text{rel}}$ | 113 s | 236 s |
| Centroid $\tilde{t}$ | 1057 s | 1117 s |

In the final step, we simulated the photo-activation on only the left half of the pattern. For this we measured the spatial intensity profile and fitted a function of the form

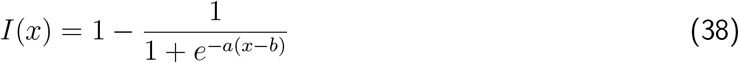

to obtain the right shape given by parameters *a* = 0.6497 and *b* = 13.186. Subsequently we modified the intensity profile such that it reaches a constant level *f* as *x* → ∞

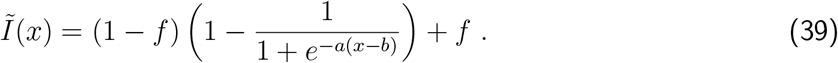

The parameter *f* ∈ [-1, 1] controls an active stress level on the non-activated side and is referred to as the *degree of active coupling*. Positive and negative values for *f* correspond to active contraction and active relaxation, respectively. The intensity profile and corresponding fit are shown in Fig. 1DS, while the activation profile *Ĩ*(*x*) for different values of *f* can be seen in Fig.4B. The time-dependent opto-stress tensor is modified by the spatial distribution of the intensity profile^1^ by multiplication

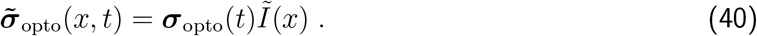

Finally, fixing all parameters as obtained by the baseline and full stimulation strain energy fits, we varied the degree of active coupling *f* as a free parameter ranging from −1 to 1 in steps of Δ*f* = 0.1, in other words, we increased the active response on the non-activated side in steps of 10%. For each value of *f*, the stress difference Δ*σ_xx_*(*x,y*) and Δ*σ_yy_*(*x,y*) between baseline and maximum strain energy were then averaged over the y-axis (Fig.4B). After that, the resulting x-profiles were normalized by integrating the right half of the curves and dividing that by the integral of the whole curve. This procedure allowed us to translate the family of curves (Fig.4B) into a relationship between the normalized stress response for *σ_xx_* and *σ_yy_* and the degree of active coupling *f* (Fig.4C).

## 2 Contour model

The observed invaginated arcs in strongly adherent cells (Fig.1A,2B) can be geometrically explained by the interplay between a surface tension *σ* associated with the contractile cortex and the resisting line tension *λ* in the strong peripheral actin bundle. In case of a homogeneous cortex one may assume the surface tension to be isotropic which yields a Laplace law predicting a constant radius of curvature *R* = *λ/σ* [11, 12]. Moreover, the observed dependence of the curvature of the arc on the spanning distance *d* of the two endpoints can be explained by assuming an elastic contribution to the line tension [11]. This modification of the simple tension model (STM) is known as the tension elasticity model (TEM) and yields a relationship *λ*(*d*) which in turn leads to an increasing *R*-*d*-relationship. However, in some cases the assumption of a homogeneous isotropic cortex fails in the presence of strongly embedded internal stress fibers. In this scenario the isotropic surface tension is modified by a directional component aligned with the direction of the internal stress fibers. This so called anisotropic tension model predicts elliptical arcs and a position dependent line tension in the fiber [13]. A comprehensive summary of the different types of existing contour models can be found in [14].

### 2.1 Anisotropic surface tension

Like all contour models, the anisotropic tension model (ATM) is based on a very general force balance equation for a slender fiber which we will motivate very briefly. The fiber is assumed to be restitant to tension only such that bending and shearing are neglected. Further we assume the fiber to start and end at discrete fixed points which resemble the focal adhesions. Each fiber has a reference shape (unstrained, stress free) and a current configuration (strained). All quantities associated with the reference shape are denoted by a Λ-symbol (Fig. 1F).

The resulting surface tension acting on the edge bundle is given by the difference of the interior and exterior stress tensors (Fig. 1E). Since the micropattern in all our experiments has two symmetry axes, we assume an anisotropic surface tension tensor of the form

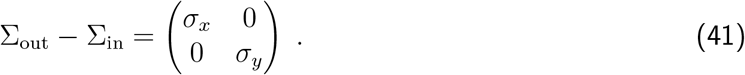

By introducing a Frenet-Serret frame as a local basis to the current configuration of the fiber

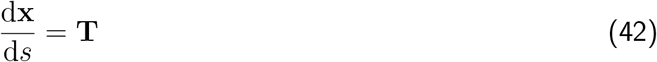

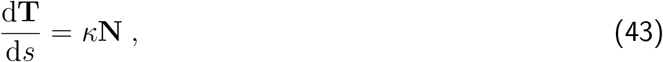

where *s* denotes the arc-length parameter along the current state, *x* the shape of the current state and *κ* the local curvature (Fig. 1F), one can derive the force balance equation by considering an infinitesimal line element in the current configuration as illustrated in Fig. 1G. For such a line element the force balance reads

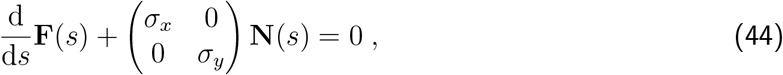

where **F**(*s*) = *λ*(*s*)**T**(*s*) always points tangential to the fiber with line tension *λ*(*s*). Finally, it can be shown that Eq. 44 leads to the equation of an ellipse

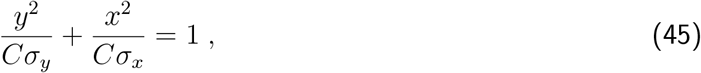

with semi-axes given by 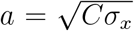 and 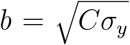. In the isotropic case, for which *σ_x_* = *σ_y_*, the ellipse attains circular shape consistent with the results of the STM and TEM.

The line tension is now a complicated function of the turning angle *θ*(*s*) given by

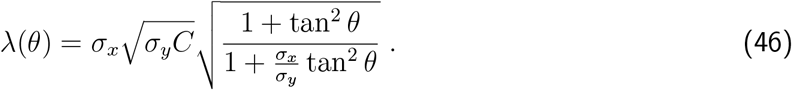

By taking derivative of this expression with respect to the turning angle *θ* one can show that the line tension has an extremum at *θ* = *θ_0_* = 0 given by

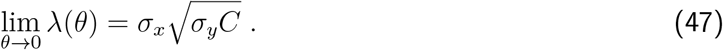

Depending on the ratio, this extremum is either a maximum for *σ_x_/σ_y_ >* 1 or a minimum for *σ_x_/σ_y_* < 1. In case of *σ_x_ = σ_y_* we obtain a constant line tension independent of the turning angle. Plots of the line tension and its derivative are shown in Fig. 2.

### 2.2 Shape analysis

Analysing cell shape is equivalent to quantifying the minimal number of key parameters like line and surface tension based on the shape of the free spanning fiber. Our goal was to apply the ATM to the TFM and fiber tracking data.

By means of our analysis we assume that all traction contribution stems from the combined action of the free spanning arc and the vertical “adherent” fiber of length *L* and add up at the intersection

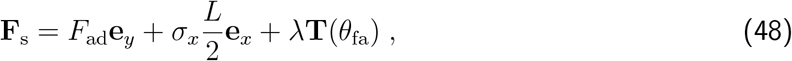

Where **F**_*s*_ is the force measured in the substrate, *θ_fa_* denotes the tangent angle at the focal adhesion and the second term is a possible contribution of the surface tension which only has an *x*-component due to the fact that the adherent fiber is straight and aligned in y-direction (Fig.2A-C). Splitting up Eq. 48 into the respective *x*-and *y*-components yields a system of two equations in the unknowns *F_ad_* and *λ*. The force **F**_s_ was obtained by dividing the traction map into four quadrants and calculating the sum for each quadrant. A similar procedure as presented in [15]. The contribution of the surface tension along the vertical fiber was estimated on TFM data as well. For the two unknowns we have

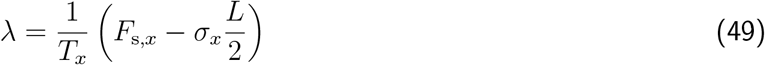

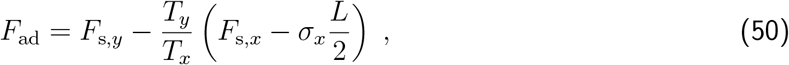

such that *λ* and *F*_ad_ can be calculated in terms of the tangent angle of the free spanning fiber at the focal adhesion.

#### 2.2.1 Ellipse shape fitting

It turned out, that fitting ellipses directly to “short” arcs is very unstable and highly depends on the initialization of the fit parameters. This is because one can find a wide range of ellipses that fit equally well. Due to large data sets of 10 to 40 cells per condition, where each cell data set consists of 60 time frames, it was not feasible to fit ellipses by hand. Therefore, we decided to use a very stable and fast circle fitting algorithm to obtain an estimate for the tangent vector at the adhesion point^2^. For the circle fitting we exploited a *Hyper least squares* algorithm presented in [16] based on algebraic distance minimization. The already determined parameters from TFM data and circle fitting are *σ_x_*, *θ_fa_*, **T**(*θ*_fa_), *λ*(*θ*_fa_). The remaining unknowns are the y-component of the surface tension tensor *σ_y_* as well as the center of the ellipse **x**_c_. Using Eq. 46 evaluated at *θ*_fa_ this yields

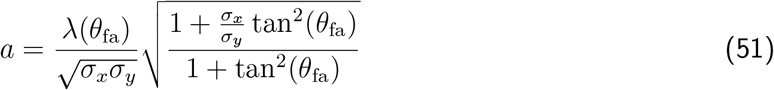

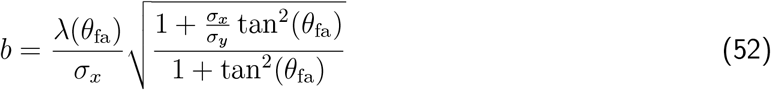

such that the shape of the ellipse purely depends on *σ_y_*. The fit was carried out by minimizing the squared distance of all tracking points along the fiber to the ellipse. The distance of those points to the ellipse was obtained by an elegant way to calculate the minimal distance of a point to the ellipse. Fig. 2C compares the standard deviations for the two fits for all conditions. In all cases, the ellipse fit yield a smaller standard deviation, although the differences vary for the different aspect ratios. The results of this analysis are summarized in Fig.2C-E.

### 2.3 Contour strain FEM-method

In order to study the effect of photo-activation on the contour and to quantify the degree of active coupling purely based on the shape of the contour we developed a discretized FEM-version of the force balance equation Eq. 44. In this context we re-formulate Eq. 44 as a function of the reference arc length parameter 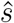 (Fig. 1F) in the reference state. The relationship between the two arc length parameters is given by stretch

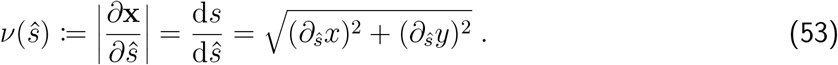

This allows to express the equation of mechanical equilibrium as

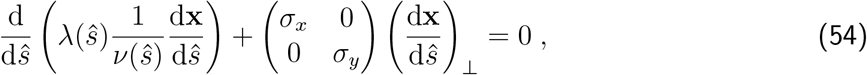

where 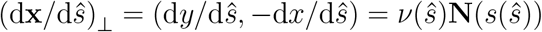. This coupled system of equations can be solved by means of a finite element implementation with mixed elements on a one-dimensional mesh. Let *ω*_1_, *ω*_2_ ∈ *D*([0, *d*]) be two test functions over the interval [0, *d*] representing the spanning distance of the unstretched straight fiber. Following the standard procedure by multiplying Eq. 54 with the test functions (one test function for each equation) and integrating it over the simulation domain yields

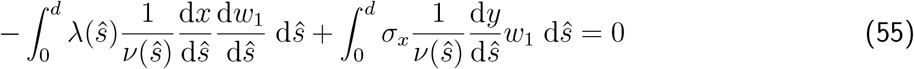

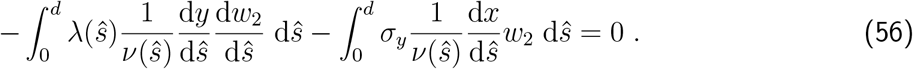

Here we used partial integration

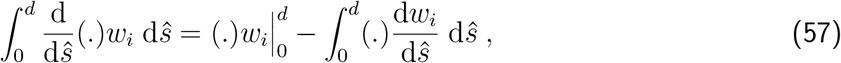

and that by construction *w_i_* = 0 on the boundary. Further we impose Dirichlet boundary conditions *x*(0) = 0, *x*(*d*) = *d*, *y*(0) = *y*(*d*) = 0 such that the endpoints of the fiber are fixed.

### 2.4 Modelling procedure for the contour finite element simulation

The modelling procedure for the contour simulation is very similar to the 2D version explained above. The aim was to quantify the active coupling between activated and non-activated part of the cell doublet. The results of the contour analysis allowed us to obtain an average ellipse by averaging the results for *a,b,σ_x_,σ_y_*. Based on actin images the spanning distance of the fiber was estimated to a value of *d* = 35 μm. An average ellipse contour was created by fixing *σ_x_* and *σ_y_* as well as the semi-axis *a*. From those values we then computed 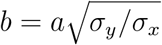. This was necessary since we averaged all those quantities independently of each other such that the averages of the single quantities not necessarily describe an elliptical arc.

In the spirit of the TEM and inspired by the work of [15] we split the line tension into an active and elastic contribution where the first accounts for the elastic properties of the cross-linking proteins within the actin bundle and the latter is an active contribution from myosin II motors such that

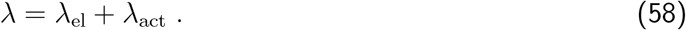

We further assumed a linear constitutive relationship between stress and strain for the elastic component

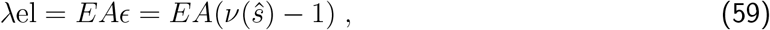

which is directly connected to the stretch as defined in Eq. 53. The rest length of the fiber is set to the spanning distance 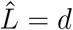. here, *EA* denotes the one-dimensional modulus of the fiber as a product of Young’s modulus *E* and the crosssectional area *A*. This value is typically around *EA* = 50 nN —350 nN [17, 15, 18]. By means of our contour simulation we set this value to *EA* = 300nN. All other fixed values for this simulation can be found in Tab. 3. Next, we minimized the simulated contour against the average contour from the contour analysis treating *λ*_act_ as a free parameter (Fig. 2D)). Subsequently, we introduced full optogenetic stimulation by defining

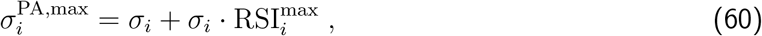

where 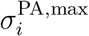 denotes the respective surface tension component at maximum strain energy, 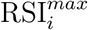 is the maximal relative surface tension increase and *i* =*x, y*. We optimized the values 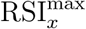, 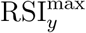 to fit the measured contour strain to the one computed with the contour FEM at maximum strain energy by additionally making sure that the values for the RSI do not exceed the from statistics experimentally obtained bounds for these values. The result of this optimization is depicted in Fig.(2E). Local photo-activation was introduced analogously to the two-dimensional case (Eq. 39) by

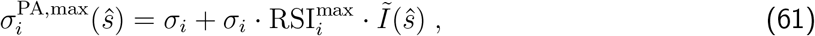

For different values of the degree of active coupling *f* we simulated the contour strain leading to the family of curves as depicted in Fig.4E. The response of the non-activated side as a function of the degree of active coupling was then obtained by the integral of the right half of the curve divided by the integral of the whole curve (Fig.4F).

**Table 3:** Fixed and opitimized parameter for the contour shape analysis by means of the contour finite element simulation.

| Parameter | Value |
| --- | --- |
| Fixed |  |
| Surface tension component $\sigma_x$ | $0.92 \text{ nN } \mu\text{m}^{-1}$ |
| Surface tension component $\sigma_y$ | $1.12 \text{ nN } \mu\text{m}^{-1}$ |
| Semi-axis $a$ | $61.94 \mu\text{m}$ |
| Semi-axis $b$ | $68.34 \mu\text{m}$ |
| One-dimensional elastic modulus $EA$ | $300 \text{ nN}$ |
| Contour fit |  |
| Active line tension $\lambda_{\text{act}}$ | $58.1 \text{ nN}$ |
| Strain fit |  |
| Relative surface tension increase $\text{RSI}_x^{\text{max}}$ | $0.11 \text{ nN } \mu\text{m}^{-1}$ |
| Relative surface tension increase $\text{RSI}_y^{\text{max}}$ | $0.24 \text{ nN } \mu\text{m}^{-1}$ |

## 3 Supplement Figures and Tables

**Figure 1:**
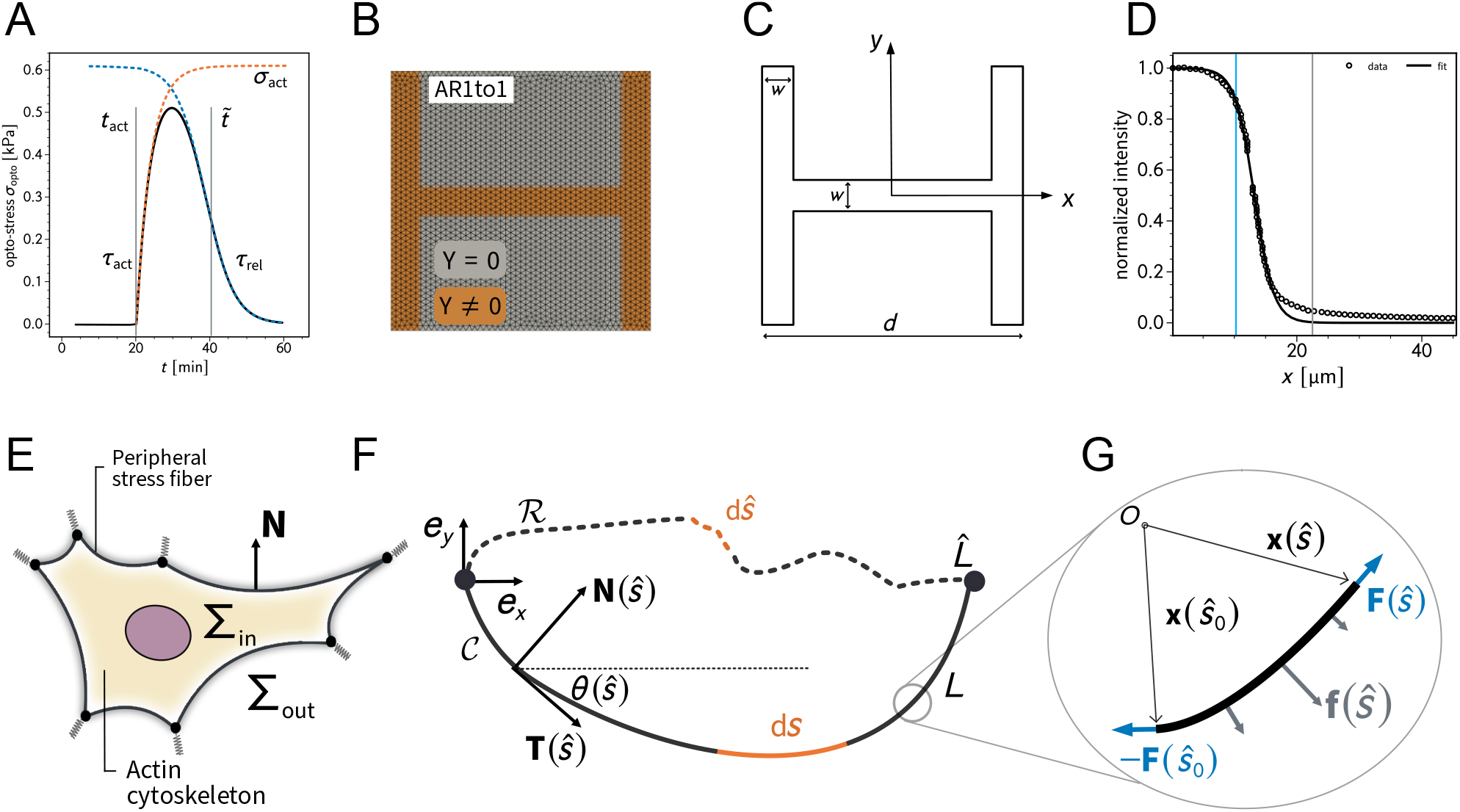
**A** shows the shape of the by optogenetic induced time dependent stress.**B** depicts the finite element mesh created with gmsh. The spring stiffness density is non-zero on the brown part of the domain. **C** is a schematic illustration of the relevant parameters to define the adhesion geometry. **D** shows the experimentally measured intensity profile of the light pulse used for photo-activation. The gray line indicates the center of the pattern (measured from left to right) while the blue line marks the inflection point of the sigmoidal fit function. **E** is a schematic illustration of the relevant quantities in the contour based description of cellular adhesion. **G,F** explain the relevant mathematical quantities to describe the equilibrium shape of a fiber subject to external loads.

**Figure 2:**
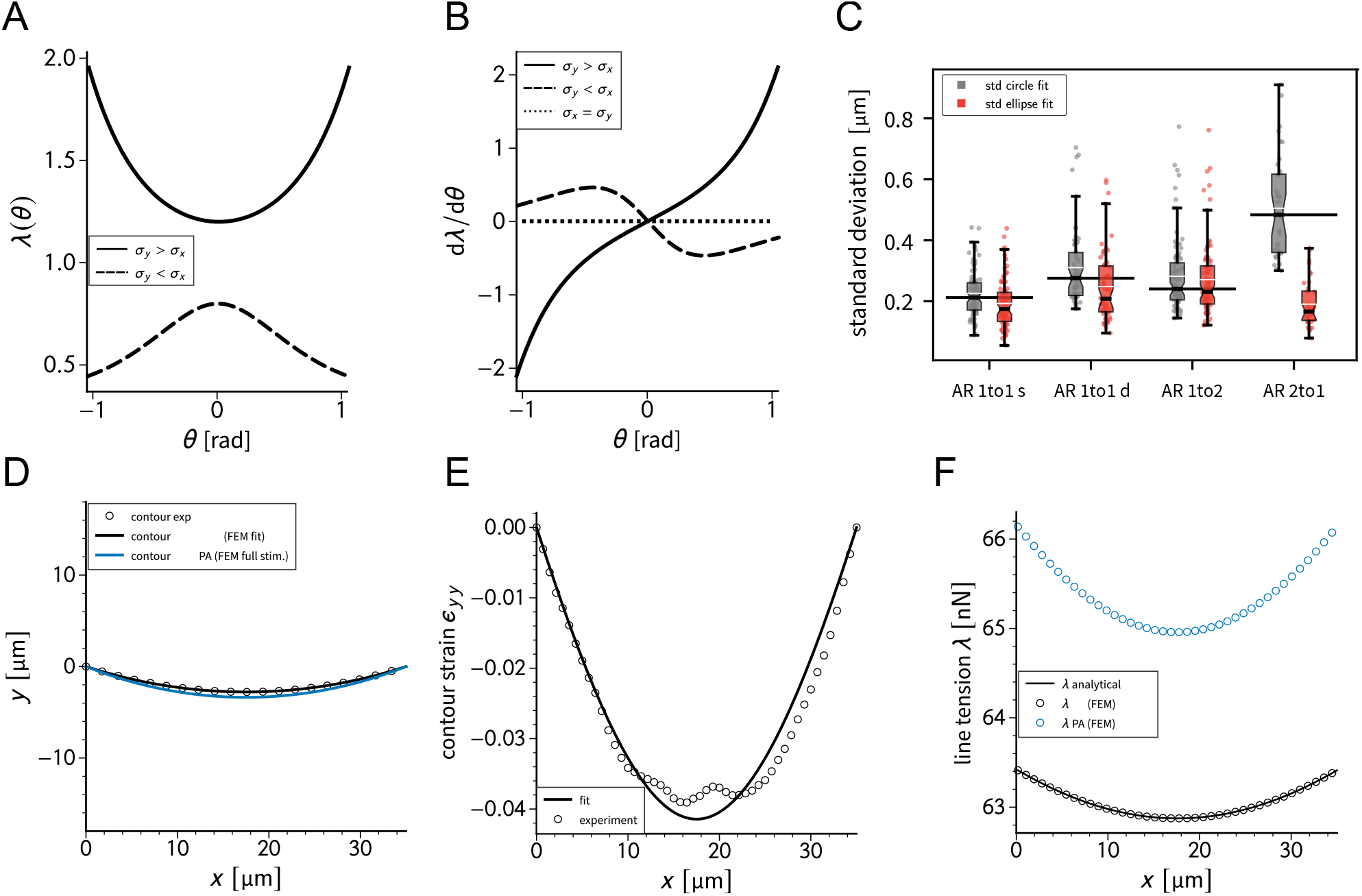
**A,B** show the predicted line tensions and the derivative with respect to the turning angle based on the analytical solution for different values of *σ_x_* and *σ_y_*. **C** compares the circle and ellipse fit of the contour of the cells for different pattern aspect ratios. **D** shows a generic cell contour for the doublet before and during photo-activation. The experimentally contour strain in *y*-direction with the respective fit from simulations and the corresponding line tensions are shown in **E** and **F**, respectively.

**Figure 3:**
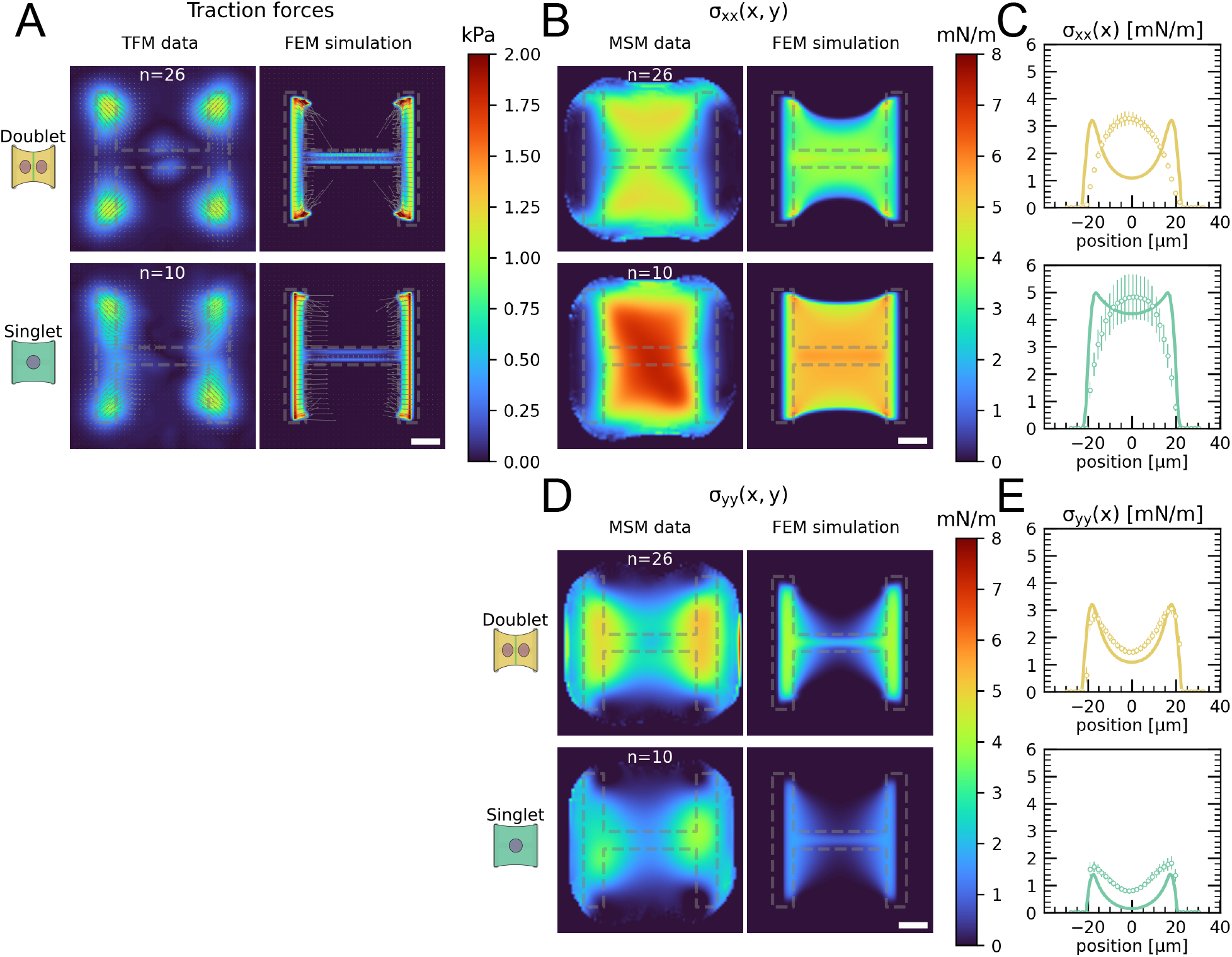
**A** Average traction stress and force maps of cell doublets (top) and singlets (bottom) on H-patterns on the left and corresponding traction stress and force maps from the FEM simulation. **B, D** Average cell stress maps of cell doublets (top) and singlets (bottom) on H-patterns on the left and corresponding cell stress maps from the FEM simulation. **C, E** Average over the y-axis of the maps in B and D. Data is shown as circles with the mean ±s.e.m., the solid line corresponds to the FEM simulations. All scale bars are 10 μm long.

**Figure 4:**
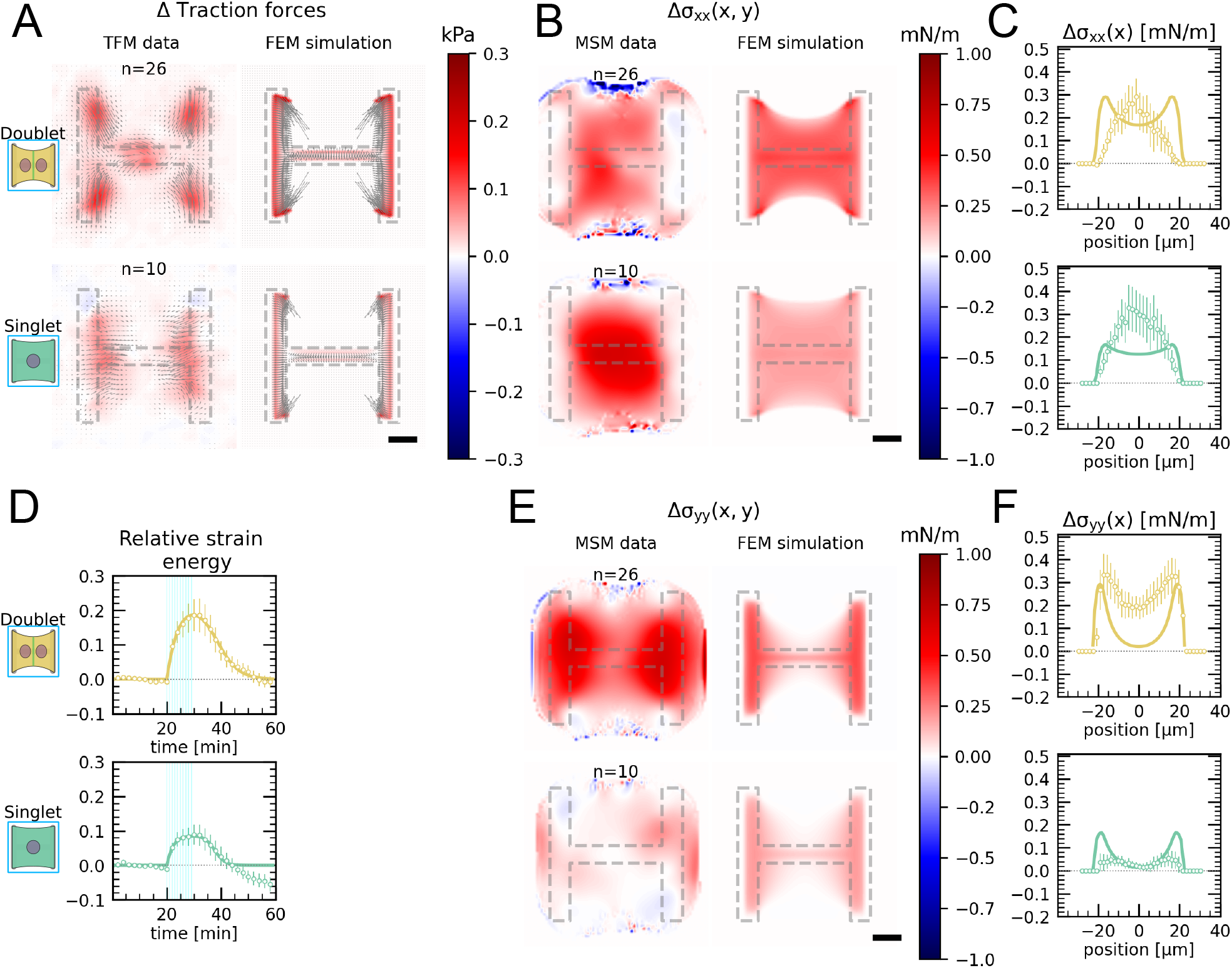
**A** Average traction stress and force map difference before and after photoactivation of cell doublets (top) and singlets (bottom) on H-patterns on the left and corresponding traction stress and force maps from the FEM simulation. **B, E** Average cell stress map difference before and after photoactivation of cell doublets (top) and singlets (bottom) on H-patterns on the left and corresponding cell stress maps from the FEM simulation. **C, F** Average over the y-axis of the maps in B and D. Data is shown as circles with the mean ±s.e.m., the solid line corresponds to the FEM simulations. **D** Relative strain energies of doublets (top) and singlets (bottom) with global photoactivation. One frame per minute was acquired for 60 minutes and cells were photoactivated with one pulse per minute for 10 minutes between minute 20 and minute 30. Strain energy curves were normalized by first substracting the individual baseline energies (average of the first 20 minutes) and then dividing by the average baseline energy of all cell doublets/singlets in the corresponding datasets. Data is shown as circles with the mean ±s.e.m and the result of an FEM simulation is shown as a solid line. All scale bars are 10 μm long.

## Footnotes

1 To keep the activation profile static in the lab-frame (eulerian frame) we incorporate the, although in many cases negligible, deformation by shifting the activation profile according to the displacement field of the previous time step such that *I(x)* = *Î*(*X* + *u_x_*). Here, the coordinate *X* is fixed in the material.

2 Although it is also possible to obtain the tangent vector directly from the fiber tracking data, we found through trial and error that this method is prone to large fluctuations.

