## Supplementary figures and images for "Force propagation between epithelial cells depends on active coupling and mechano-structural polarization"

### Figure S1

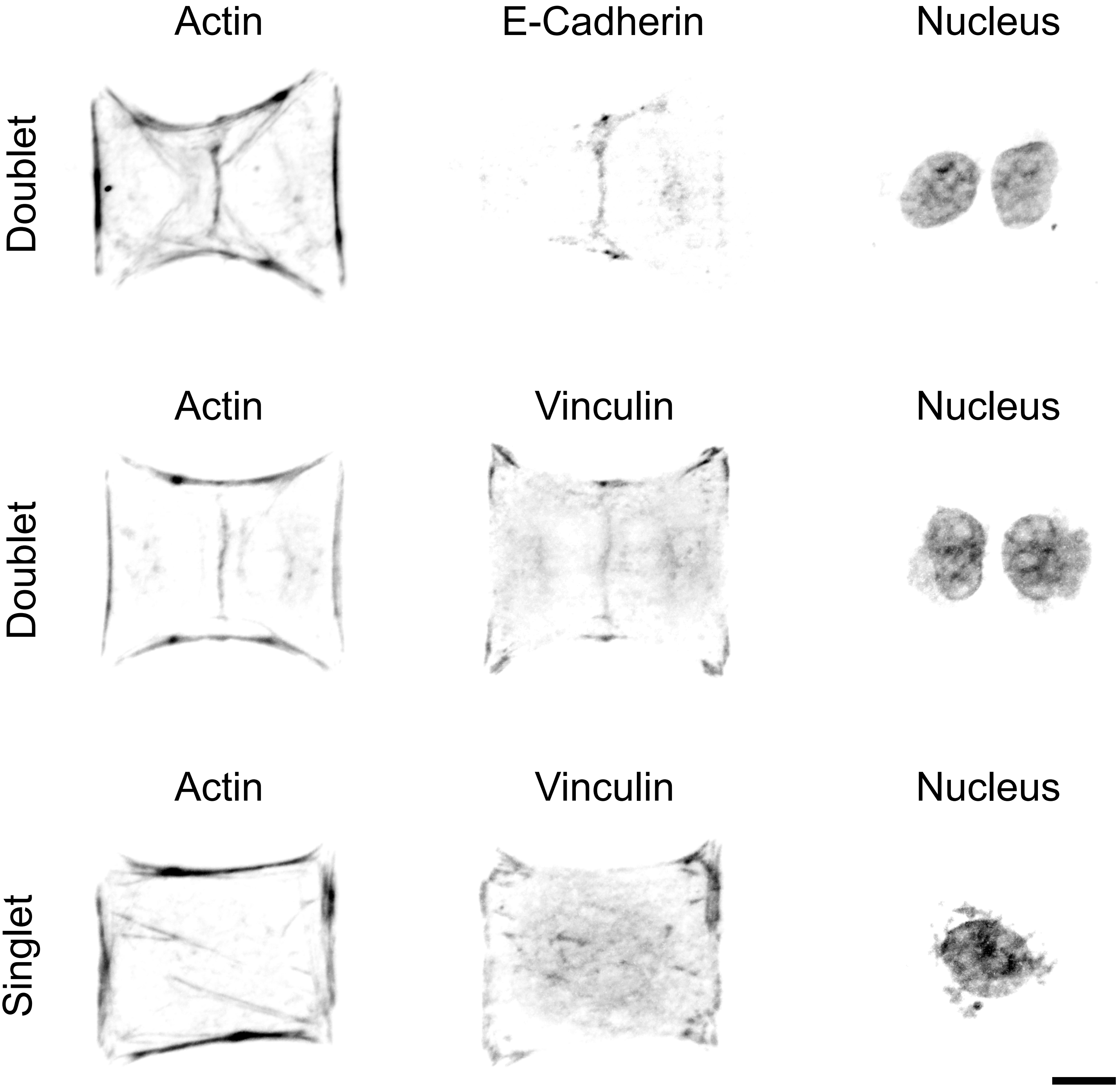

### Figure S2

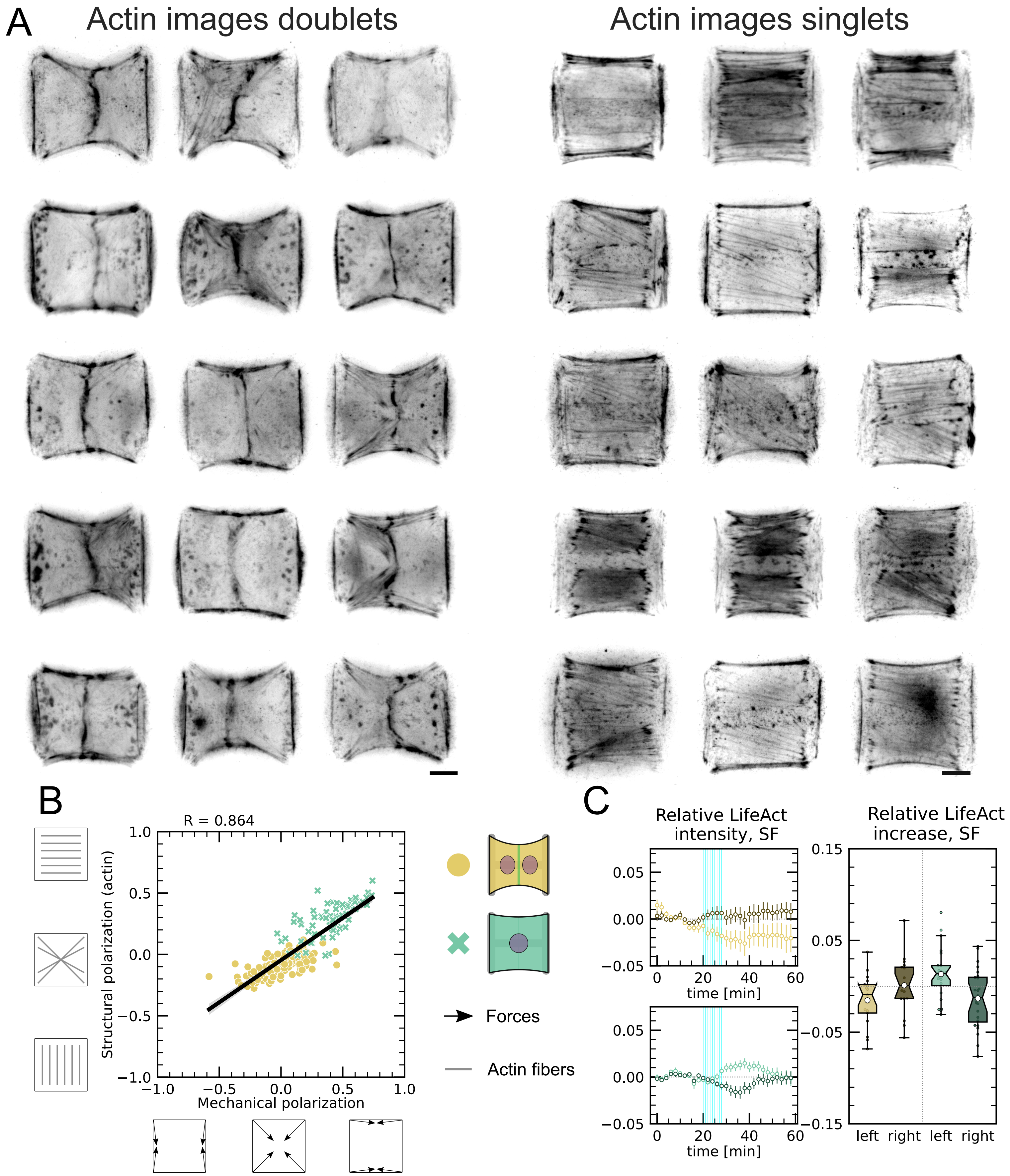

### Figure S3

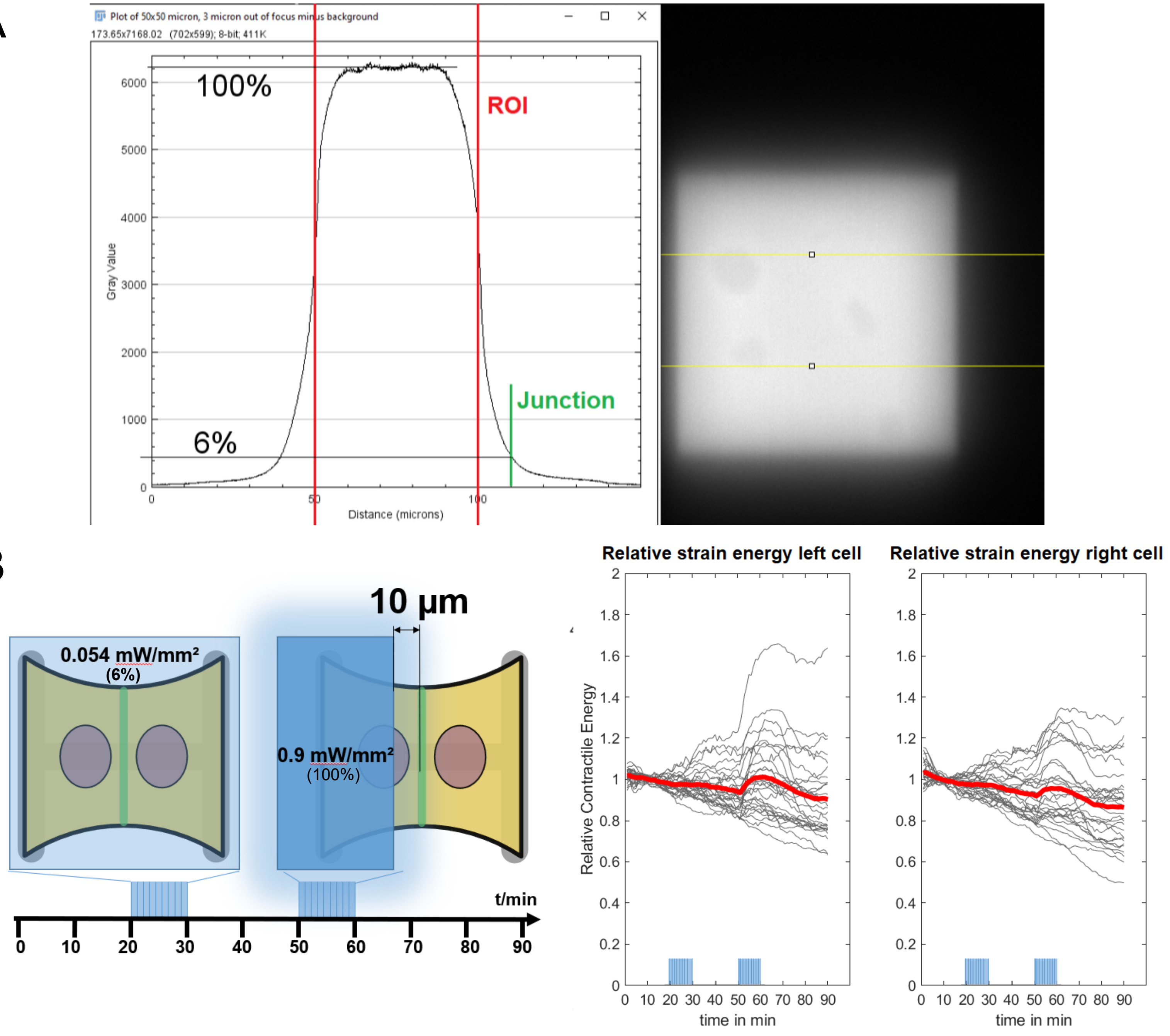

### Figure S4

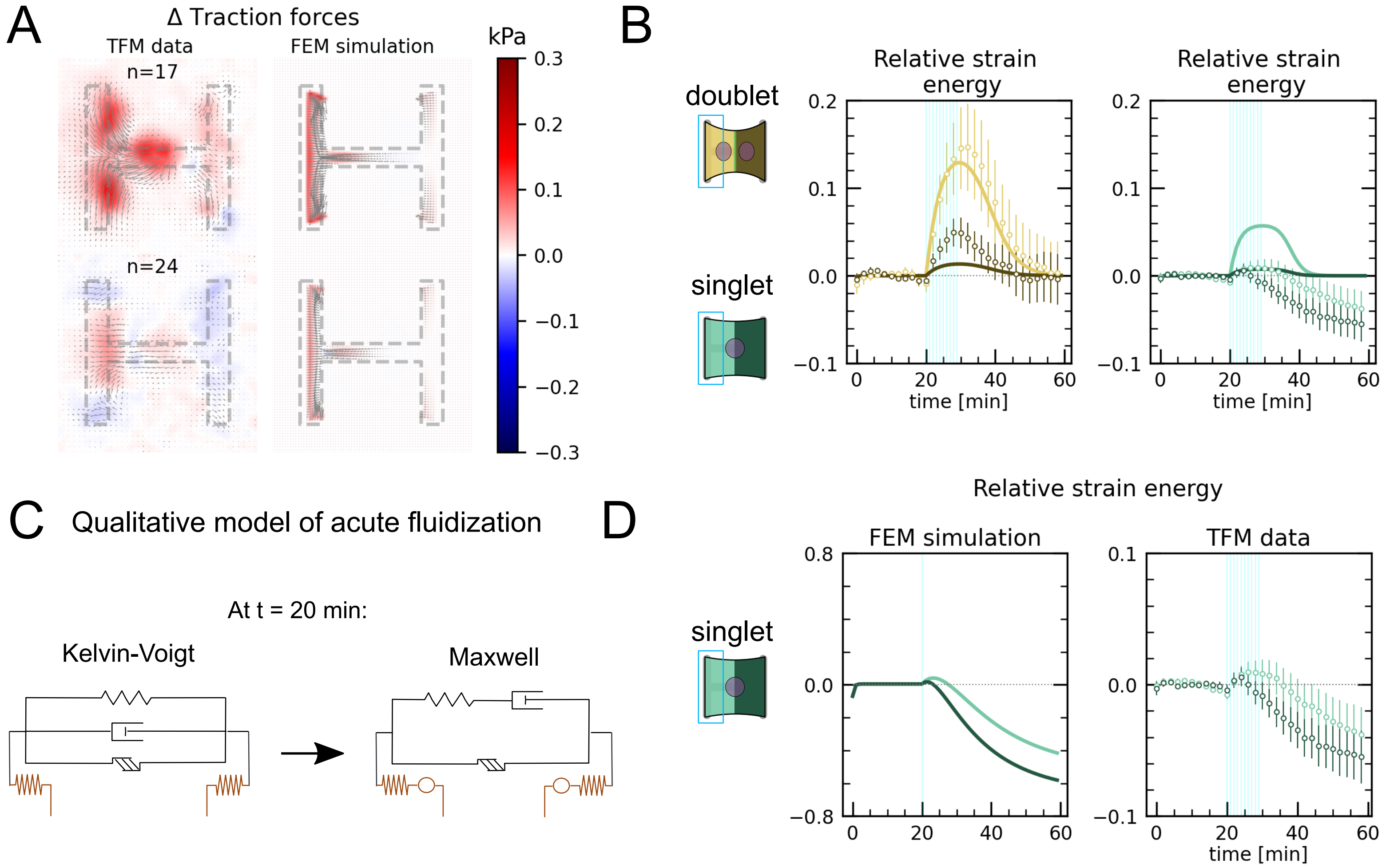

### Figure S5

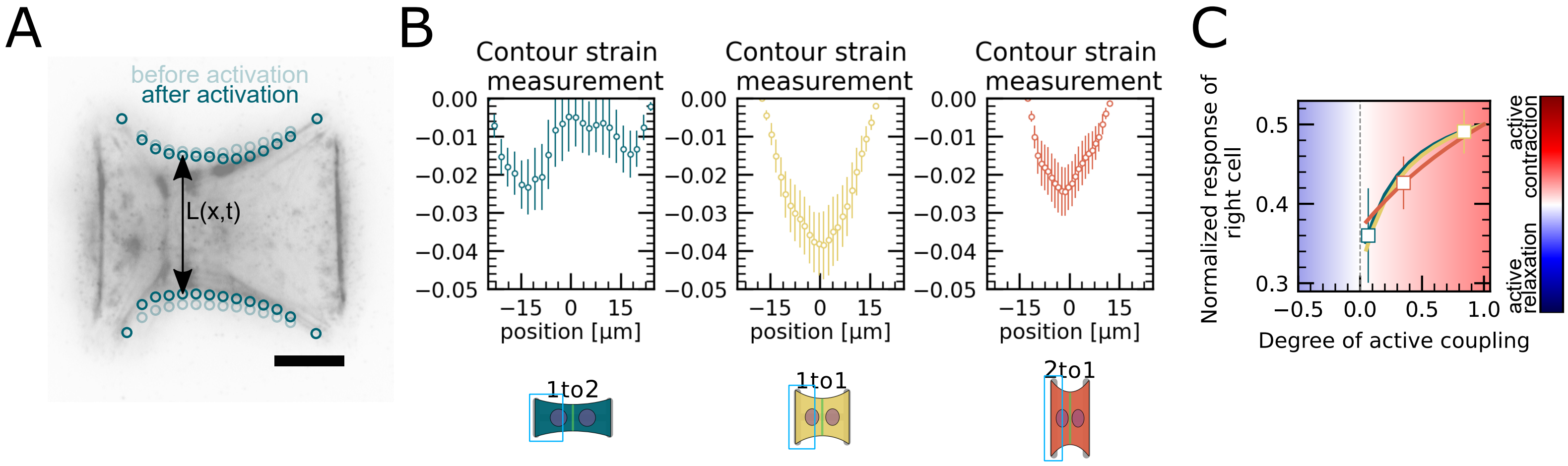

### Figure S6

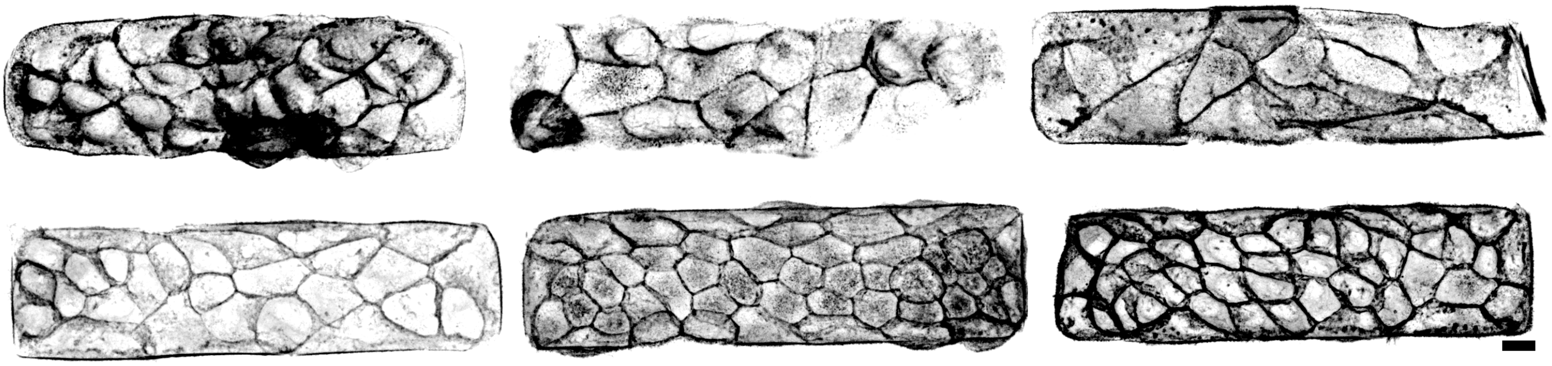

### Movie S1

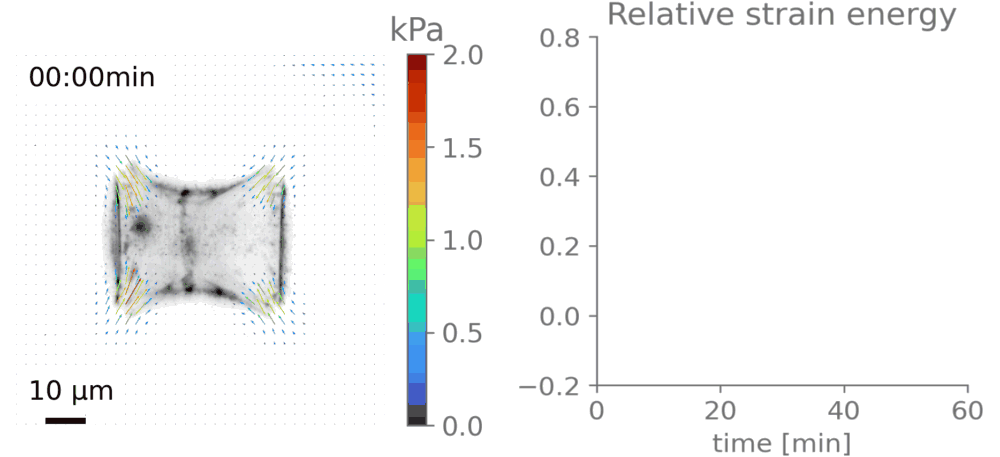

### Movie S2

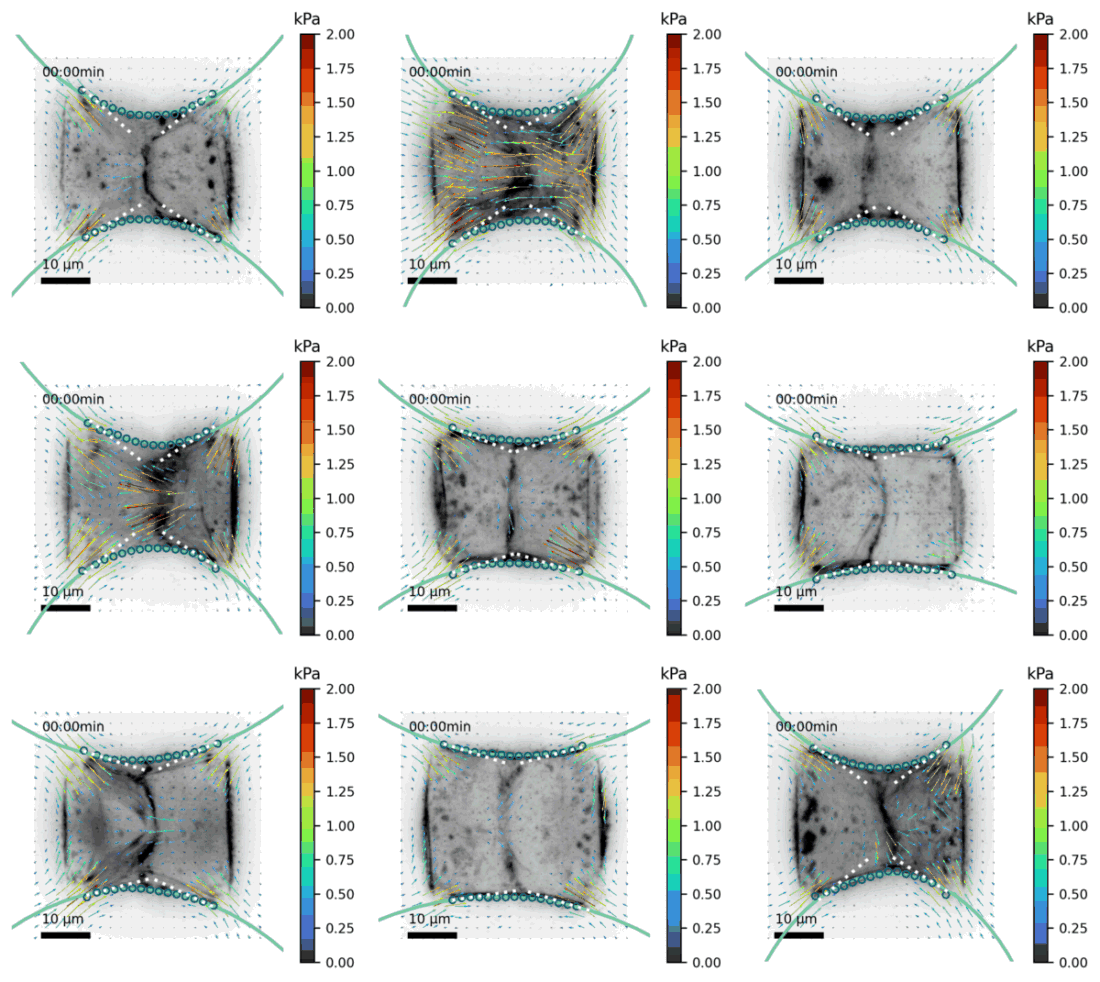

### Movie S3

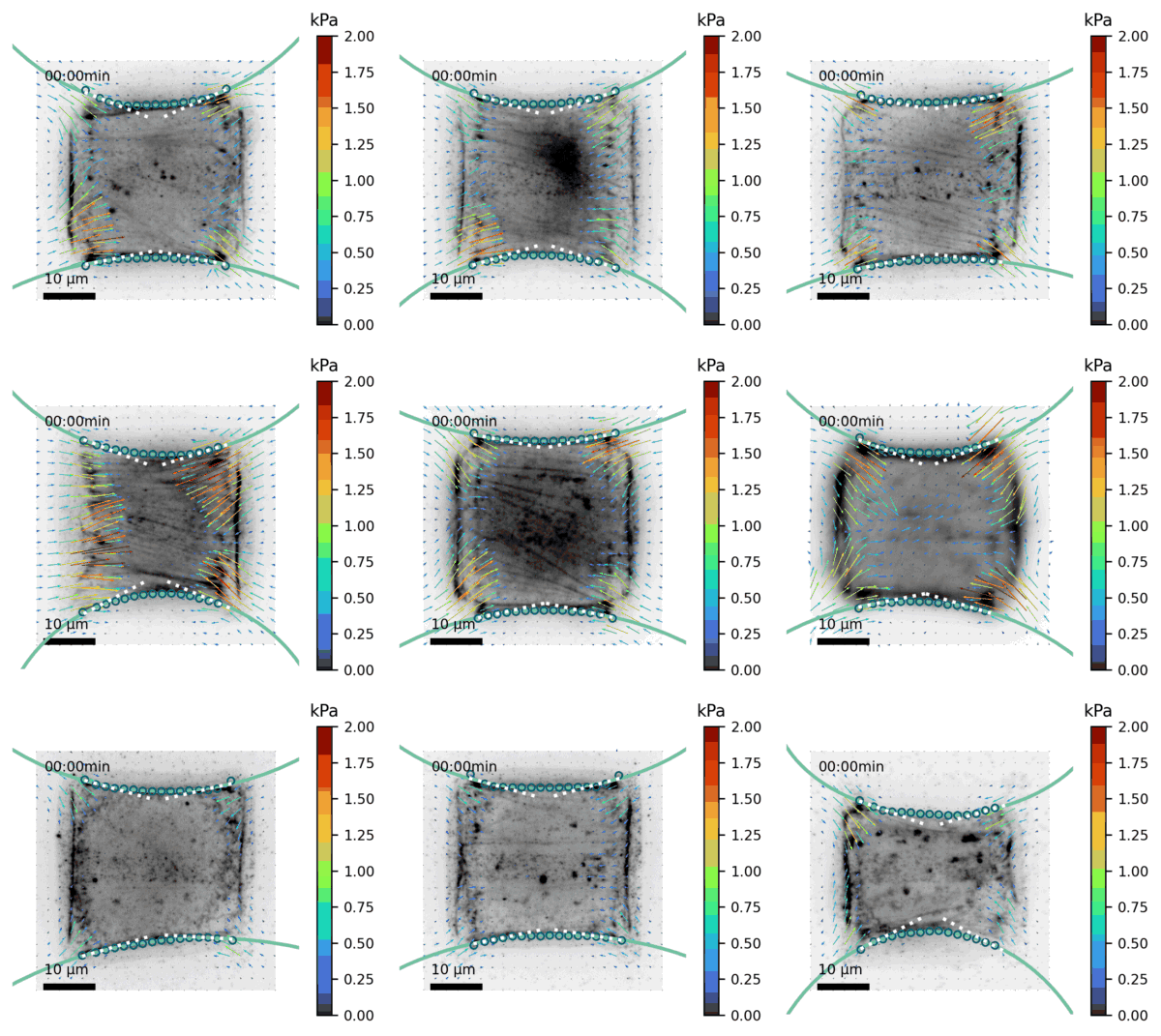

### Movie S4

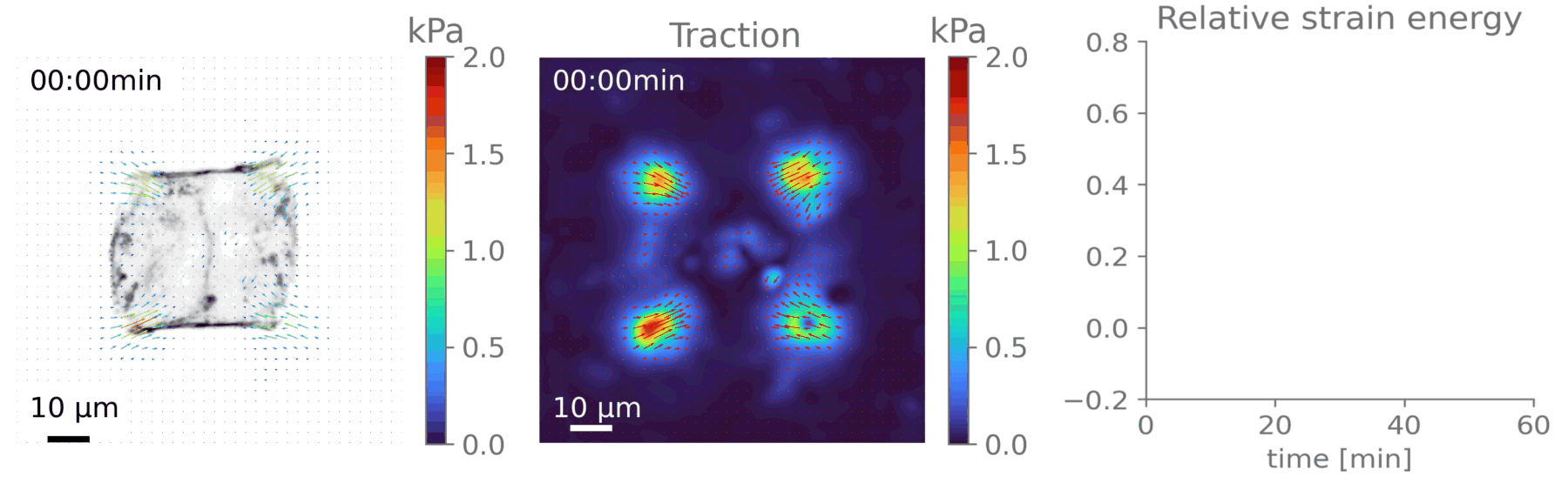

### Movie S5

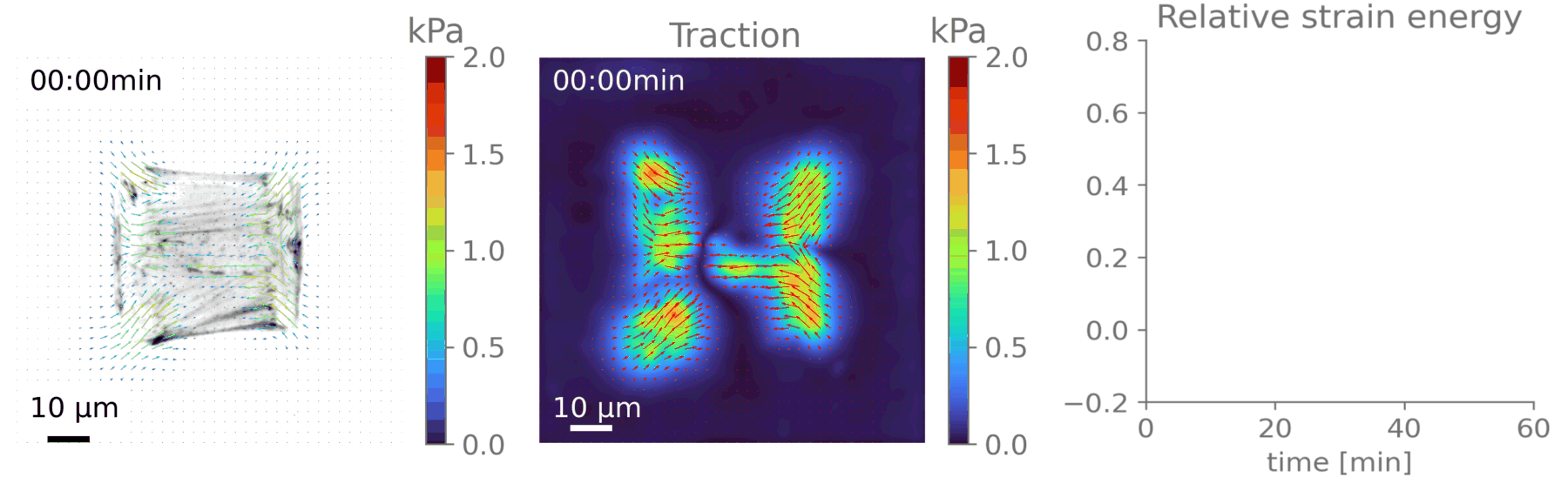

### Movie S6

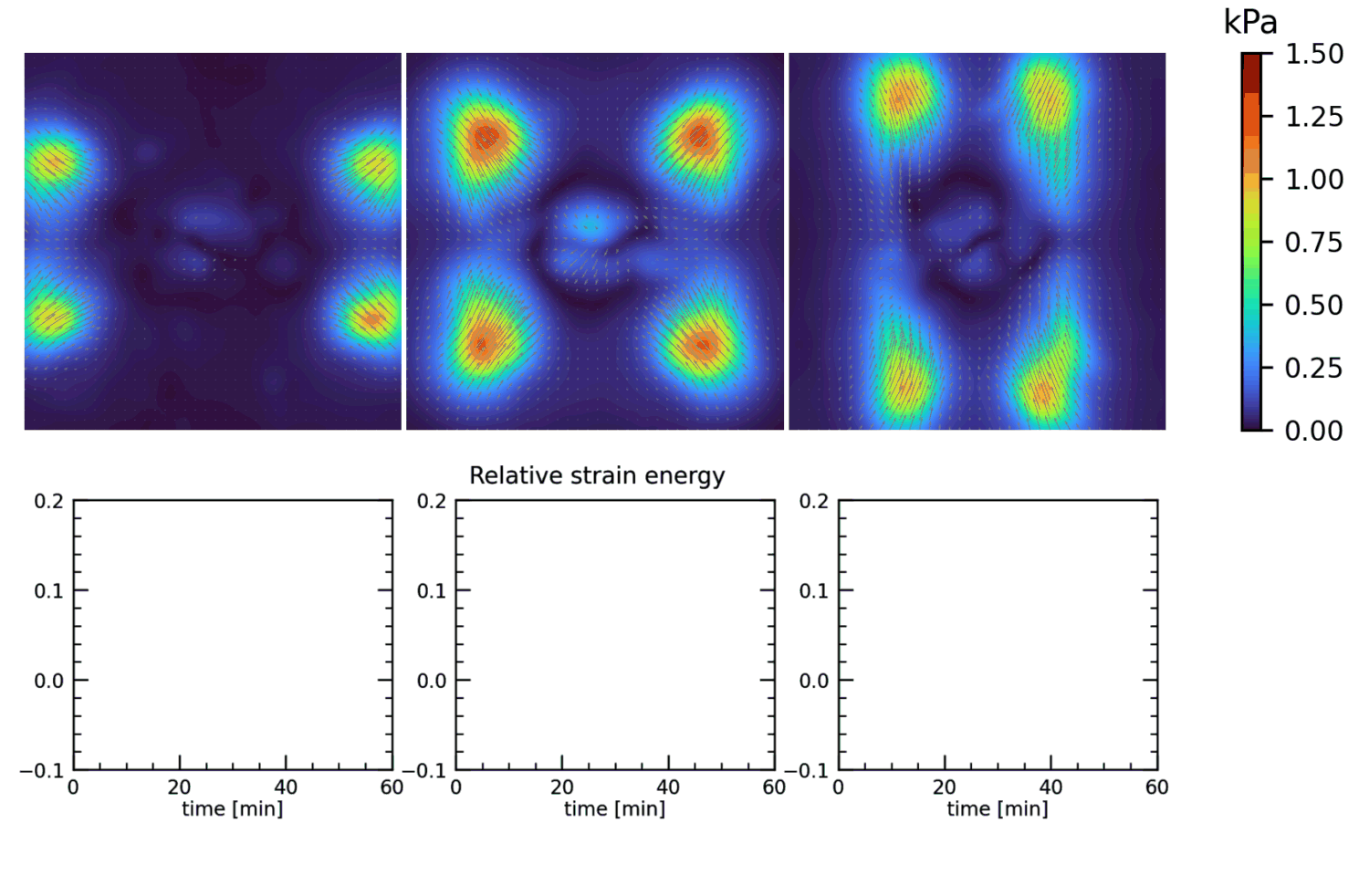

### Movie S7

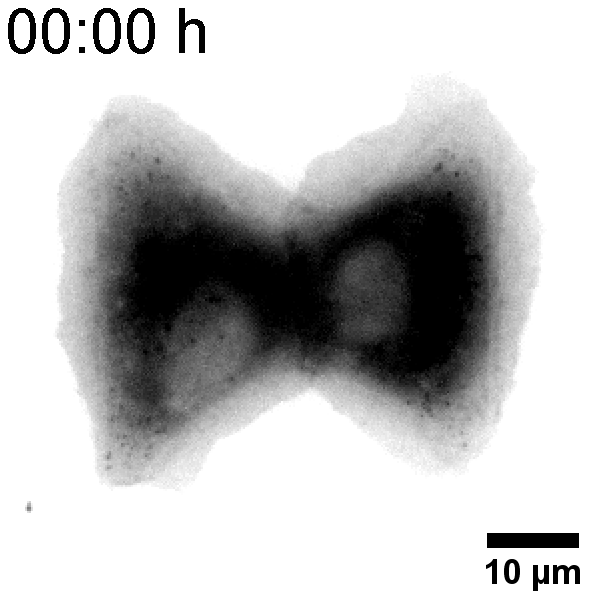
